# Correction of age-associated defects in dendritic cell functions enables CD4^+^ T cells to eradicate tumors in the elderly

**DOI:** 10.1101/2022.11.08.515695

**Authors:** Dania Zhivaki, Francesco Boriello, Stephanie N. Kennedy, Charles L. Evavold, Kristin M. Bahleda, Kate L. Chapman, Ivan Zanoni, Jonathan C. Kagan

**Affiliations:** Harvard Medical School and Division of Gastroenterology, Boston Children’s Hospital, Boston, Massachusetts, USA; Current Employer, Corner Therapeutics, Watertown, Massachusetts, USA; Harvard Medical School and Division of Immunology, Boston Children’s Hospital, Boston, Massachusetts, USA; Current Employer, Generate Biomedicines, Cambridge, Massachusetts, USA

## Abstract

Defective host defenses later in life are associated with changes in immune system activity. The means to correct immune defects to ensure immunity in the elderly are undefined. In this study, we found that CD8^+^ T cells, which are necessary for anti-tumor immunity in young mice, are not required to eradicate the same cancers later in life. Rather, CD4^+^ T cells drive anti-tumor immunity in elderly mice. The generation of anti-tumor CD4^+^ T cells requires multiple dendritic cell (DC) activities that are elicited by immune agonists known as hyperactivators, but not by adjuvants that model those used clinically. DC hyperactivators correct age-associated defects in DC migration and T cell co-stimulation while enabling NLRP3 inflammasome activities within living cells. These combined activities enable DCs to induce TH1-skewed T cells that persist into old age and eliminate implantable tumors. These results raise the possibility of correcting age-associated immune defects through DC manipulation.

## Introduction

The mammalian immune system is a dynamic entity, which undergoes radical changes in cellular composition and function based on life experiences. One such life experience is the process of aging, where decreased immune system activities lead to an increased risk of infection and cancer^1^. Elderly humans display a diminished abundance of naïve T cells, as compared to young adults, resulting in defective generation of *de novo* T cell responses to newly encountered antigens. Aging is also associated with alternations in dendritic cells (DCs), which in young adults are robust stimulators of protective T cell responses. Age-associated changes in DC functions include a striking deficiency in migratory activities and the reduced production of select cytokines, both of which are necessary for the stimulation of protective T cell responses^2–7^. These defects in immune cell function, along with others, may underlie the lack of effective host defenses observed in elderly populations^8^.

While several age-associated immune cell defects have been documented, few examples of correction of these defects exist. Indeed, prophylactic immunostimulants (*i.e.* vaccines) are unable to correct immune defects in the elderly and display diminished efficacy as the age of the patient population increases^8–10^. Similarly, therapeutic immune checkpoint inhibitors, such as those targeting programmed cell death 1 (PD-1), programmed cell death ligand 1 (PD-L1) and cytotoxic T lymphocyte antigen 4 (CTLA-4) pathways, yield variable safety and efficacy outcomes in very young or old cancer patients, as compared to young adult cohorts of patients^11–13^. Current immunotherapies and vaccination approaches often overlook the alteration of the ageing immune system and are rather designed to treat young and healthy adults^8^.

Using our knowledge of the young adult immune system as a guide, several principles of immunity have been established. Central to immune system functions are the DCs, which survey the tissues of mammals for molecular indicators of threats to the host. Common molecular indicators include microbial products and host-derived indicators of tissue injury, which stimulate Toll-like Receptors (TLRs) or other pattern recognition receptors (PRRs) on DCs^14^. Upon PRR activation, three DC activities are induced that instruct T cell mediated immunity. These activities include 1) MHC-mediated presentation of antigens by DCs to naïve T cells, 2) upregulation of co-stimulatory molecules such as CD40, CD80 and CD86^15^, which ensure robust DC interactions with naïve T cell in the lymph node, and 3) secretion of cytokines that differentiate naïve T cells into effector cell subsets. A prime example of the latter is the production of IL-12p70 by TLR-stimulated DCs, which prompts CD4^+^ T cell differentiation into type 1 T helper (TH1) cells^16^.

In addition to these three DC activities, DCs must acquire the ability to migrate to the lymph nodes that drain the tissue where the threat to the host was encountered^17^. These nodes contain abundant pools of naïve T cells and are the primary sites of DC-T cell interactions that stimulate new adaptive immune responses. Finally, studies in recent years have underscored the importance of the cytokine interleukin 1 (IL-1) in T cell functions^18^. IL-1β can act directly on T cells via the IL-1 receptor to enhance T cell polyfunctionality, antigen-specific T cell expansion, the differentiation of naïve T cells into memory cells ^19–23^ and the reactivation of memory T cells^24^. Consistent with this idea, studies from Paul and colleagues demonstrated that addition of recombinant IL-1β to immunogens massively enhanced the production of antigen-specific CD4^+^ and CD8^+^ T cell responses in mice^21, 22^.

Despite the ability of exogenous IL-1β to potentiate T cell responses, approaches to stimulate IL-1β production from living DCs are not commonly used to drive protective immunity in mice or the clinic. For example, adjuvants that stimulate TLRs induce the expression of the gene encoding pro-IL-1β, but these pathways do not stimulate the primary drivers of bioactive IL-1β release from DCs—inflammasomes^25, 26^. As such, TLR stimulated DCs do not release IL-1β into the extracellular space. Conversely, aluminum hydroxide (alum), the most-commonly used FDA-approved adjuvant for clinical use, is an inflammasome agonist that drives IL-1β production from stimulated cells. However, alum induces the death of DCs via pyroptosis, which should undermine other DC activities that are important for T cell instruction ^27–29^. The ideal strategy of DC stimulation may be one that stimulates robust DCs migration to the dLN and induces IL-1β production while maintaining viability. Recent studies have revealed such a state of DC activation, which has been dubbed hyperactivation^30^.

Hyperactive DCs are elicited by the exposure of these cells to TLR ligands and an oxidized phospholipid known as PGPC. The former is a molecular indicator of microbial encounters whereas the latter is an indicator of tissue injury. TLR ligands alone or TLR ligands+PGPC stimulate antigen presentation and costimulatory molecule expression by DCs, as well as effector T cell differentiating cytokine expression. However, while TLR ligands do not induce robust DC migration to lymph nodes and do not stimulate IL-1β production by DCs, TLR ligands+PGPC induce both of these activities. The addition of these two activities to the immunostimulatory repertoire renders DCs not merely active (as TLR stimulated cells are often dubbed), but rather hyperactive^30^. Hyperactive DCs have a superior ability to stimulate anti-tumor CD8^+^ T cell responses^30^, as compared to DCs stimulated with classical activation stimuli (*e.g.* TLR agonists or alum). Despite this knowledge, the functions of hyperactive DCs in most contexts of immunity are undefined, in particular in the context of age-dependent immunity. As TLR-stimulated DCs are incapable of correcting defects in T cell functions in the elderly^2, 31^, we considered the possibility that hyperactivators of DCs may yield enhanced protective function in the elderly.

In this study, we found that elderly mice are less able to survive challenges with implantable tumors than young mice. This defect in protective immunity cannot be corrected by T cell targeting checkpoint inhibitor therapies (*e.g.* anti-PD-1 or anti-CTLA4), but can be corrected by hyperactive DCs. These cells use tumor antigens to generate new anti-tumor T cell responses in an age-agnostic manner. We identified a shift in the means by which hyperactive DCs elicit anti-tumor immunity in young and elderly mice. Whereas CD8^+^ T cells mediate tumor rejection in young mice, the same tumors are controlled by CD4^+^ T cells in elderly mice. We demonstrate that hyperactive DCs can elicit protective antigen-specific CD4^+^ T cell responses over a year after immunization, or upon primary immunization of elderly mice. Other DC activation states are unable to induce T cell immunity in the elderly. Mechanistically, we found that NLRP3 inflammasomes produce IL-1β from hyperactive DCs to accelerate CD4^+^ T cell differentiation into a TH1-skewed response. These responses, in elderly mice and in primary human cells, are required for anti-tumor immunity. Immune defects in the elderly are therefore not insurmountable, suggesting that age-dependent tailoring of immunostimulatory activities may ensure life-long immunity.

## Results

### DC hyperactivators correct defective T cell mediated anti-tumor immunity in elderly mice

We began our studies with the observation that elderly C57BL/6J and Balb/cJ mice exhibit poor anti-tumor immunity as compared to young mice. Eight weeks (young) or 68 weeks (elderly) mice were subcutaneously (s.c.) injected with B16 melanoma tumor cells that express the model antigen ovalbumin (OVA), which are referred to as B16OVA tumor cells, or with CT26 colon adenocarcinoma tumor cells that do not express OVA. Despite being implanted with equivalent numbers of tumor cells, elderly mice were less capable than young mice in restricting tumor growth and died faster than their younger counterparts **(Figure 1A)**. To determine if mouse models of clinical immunotherapies could correct age-associated defects in anti-tumor immunity, we treated mice bearing 2-3mm tumors with antibodies that block PD-1 (anti-PD-1) or CTLA4 (anti-CTLA4). Anti-PD-1 treatments of young mice harboring B16OVA tumors were protective, resulting in tumor rejection in 100% of mice **(Figure 1B)**. Surprisingly, anti-PD-1 was ineffective as a therapy in old mice **(Figure 1B)**. Similar trends were observed in Balb/cJ mice implanted with CT26 tumors, where the protective effects of anti-PD-1 were only observed in young mice **(Figure S1A)**. We note that even in young mice, the CT26 model was less responsive to anti-PD-1 treatments than the B16OVA model. These findings are consistent with prior work in CT26^32^. Similar findings were made when mice bearing B16OVA tumors were treated with anti-PD1 alone, anti-CTLA4 alone or anti-PD1+anti-CTLA4 **(Figure S1B)**. While these checkpoint inhibitor treatments induced protection in young mice, the same treatment had some effect in delaying tumor growth in old mice, but all mice in these elderly cohorts died **(Figure 1B-S1B)**.

**Figure 1.**
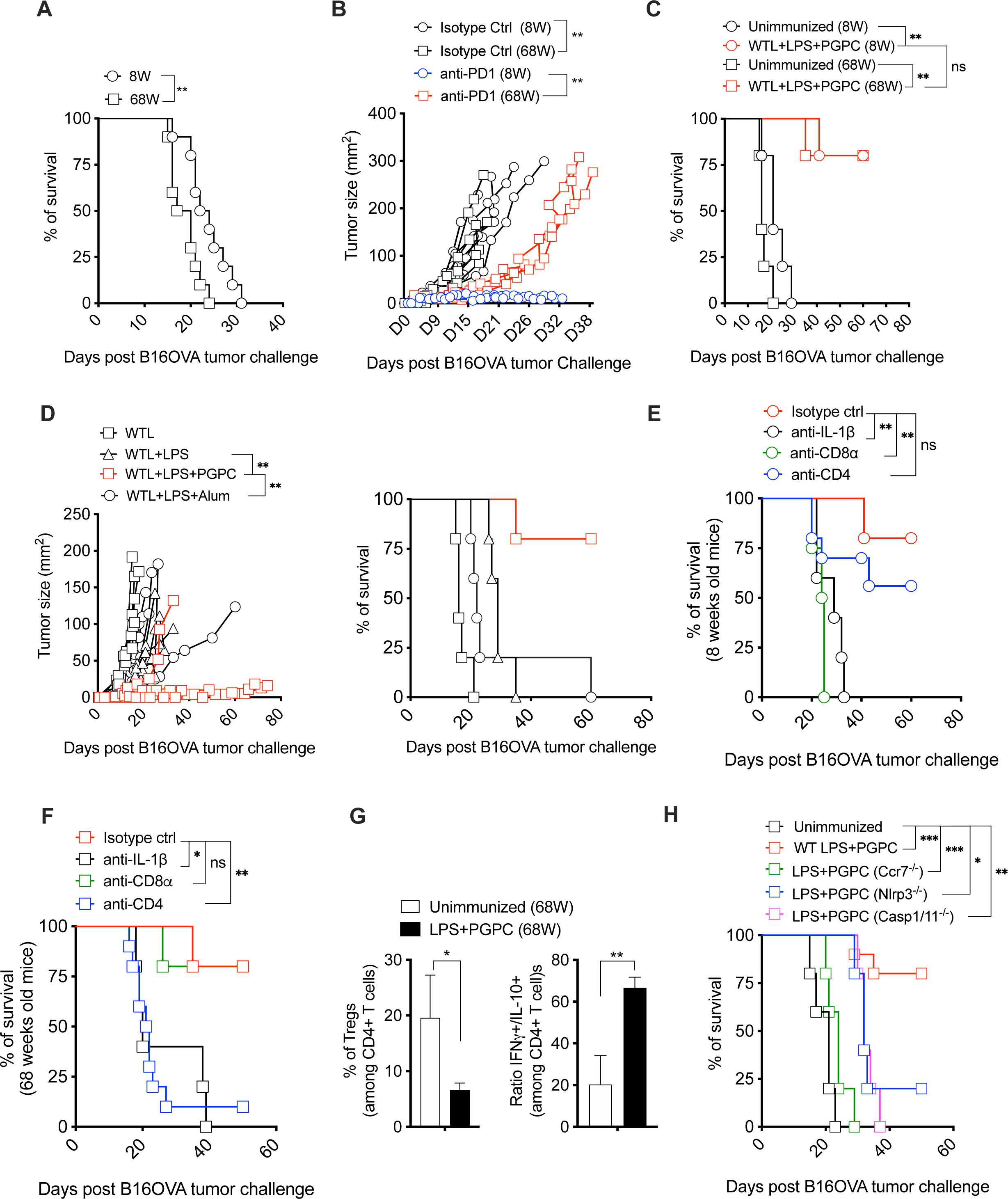
CD4^+^ T cells mediate anti-tumor immunity in young and old mice. (A-D) 8 weeks or 68 weeks old C57BL/6J mice were injected subcutaneously (s.c.) on the right flank with B16OVA cells. When tumors reached 2-3 mm of size, mice were either (A) left untreated (unimmunized) or (B) treated i.p. with anti-PD1 antibodies alone, (A) The percentage of mice survival was monitored every 2 days (n=10 mice per group). (B) Tumor size in mm^2^ was measured every two-three days (n=3 mice per group). (C) B16OVA tumor-bearing mice were immunized with B16OVA whole tumor lysate (WTL) and LPS plus PGPC emulsified in IFA. Tumor size in mm^2^ was measured every two-three days (n=5 mice per group). (D) 68 weeks old C57BL/6J mice were injected s.c. on the right flank with B16OVA cells. When tumors reached 2 mm, mice were immunized on the left flank with PBS alone or with and LPS, or with LPS plus PGPC all emulsified in IFA, or mice were immunized with LPS plus Alum. Tumor size in mm^2^ was measured every two-three days (Left panel). The percentage of mice survival was monitored every 2 days (Right panel). (E) 8 weeks old or (F) 68 weeks old C57BL/6J mice were injected s.c. on the right flank with B16OVA cells. When tumors reached 2mm in size, mice were either left unimmunized or were immunized with LPS plus PGPC with or without neutralizing anti-CD4, or anti-IL-1β or anti-CD8a antibodies. Percentage of mice survival was monitored every 2 days (n=5-10 mice per group). (G) 68 weeks old mice were injected s.c. on the right flank with B16OVA cells. When tumors reached 2-3mm in size, mice were either left unimmunized or were immunized with LPS plus PGPC. 15 days post immunization, CD45 TILs were enriched using anti-CD45 microbeads. The percentage of Tregs as CD4^+^ Foxp3^+^ T cells was measured by flow cytometry (Upper panel). CD45 TILs were activated for 24 hours with anti-CD3/CD28, then treated with PMA and ionomycin for 5 hours. The ratio of IFNg over IL-10 producing CD4^+^ T cells in the tumor microenvironment was measured by intracellular staining. (H) 68 weeks old wild type or *Ccr7*^-/-^ or *Casp1/11*^-/-^ or *Nlrp3*^-/-^ mice were injected s.c. on the right flank with B16OVA cells. When tumors reached 2-3mm in size, mice were either left unimmunized or were immunized with WTL and LPS plus PGPC. Percentage of mice survival was monitored every 2 days (n=5 mice per group).

PD-1 antibodies are thought to induce protective immunity in several ways, with the best described mechanism being to invigorate the inflammatory activities of T cells^33^. To determine if age-dependent defects in anti-tumor immunity could be corrected by DC-targeting approaches, we examined the ability of distinct DC agonists to stimulate protective immunity in young and elderly mice. Eight week or 68 week old mice were implanted s.c. with B16OVA tumor cells. When tumors reached 2-3mm in size, mice were either left unimmunized, or they were immunized s.c. with DC hyperactivators consisting of LPS and PGPC, along with whole tumor lysates (WTL) from B16OVA cells as an antigen source. Alternatively, mice were immunized with LPS+WTL, or LPS+alum+WTL. The latter stimulus represents a model of the FDA-approved adjuvant alum, which is used clinically on its own or in combination with LPS-like TLR agonists^34, 35^. The immunization of tumor-bearing mice with LPS+PGPC+WTL induced tumor rejection in ∼80% of all mice, regardless of age **(Figure 1C)**. In contrast, anti-tumor immunity was ineffective when elderly mice were immunized with WTL in combination with LPS alone or with LPS+alum **(Figure 1D).** These results indicate that unlike anti-PD-1 treatments or other strategies of DC stimulation, DC hyperactivators can correct age-associated defects in host defense.

Hyperactive DCs and PD-1 antibodies induce anti-tumor immunity via the actions of T cells in young mice^30, 36–38^, yet only treatments that stimulated the former provided tumor protection in elderly mice **(Figure 1B-D)**. These results raised the possibility that the mechanisms of protective immunity may differ in an age-dependent manner. To address this possibility, we first assessed the abundance of CD4^+^ and CD8^+^ T cells among TCRα^+^ splenic cells in 8 and 68 weeks old mice. We found that 68 weeks old mice contained four-fold less naïve T cells than young mice **(Figure S1C-D).** In addition, elderly mice displayed a substantial reduction in CD8^+^ T cell abundance, as compared to young mice **(Figure S1C-D)**. Similar defects in T cell abundance have been reported in elderly humans^31, 39^ or aged mice. Based on the decreased abundance of CD8^+^ T cells in elderly mice and the decrease in overall naïve T cells in these mice, we examined the requirement of CD4^+^ and CD8^+^ T cells in anti-tumor immunity in each age group.

CD4^+^ or CD8α^+^ T cells were depleted from mice containing 2-3 mm B16OVA tumors using neutralizing antibodies. These neutralizing antibodies were injected intraperitoneally (i.p.) one day prior to the day of therapeutic immunization with LPS+PGPC+WTL. Five subsequent injections of anti-CD4 and anti-CD8α antibodies were performed every 2 days following therapeutic immunization. LPS+PGPC+WTL injections induced tumor rejection that was dependent on CD8a^+^ T cells in young mice **(Figure 1E)**, a finding consistent with prior work^30^. Surprisingly, CD8α^+^ T cell depletion did not disrupt anti-tumor responses in elderly mice. Rather, the depletion of CD4^+^ T cells eliminated anti-tumor responses induced in elderly mice **(Figure 1F)**. These age-associated changes in T cell dependence for host defense were also observed in Balb/cJ mice implanted with CT26 tumors **(Figure S1E).** In this latter model, the immunization of mice with LPS+PGPC+WTL led to tumor rejection in 100% of mice. The anti-tumor protection in elderly mice was dependent on CD4^+^ T cells and IL-1β, since the immunization of mice in the presence of CD4 or IL-1β antibodies prevented tumor rejection **(Figure S1E)**.

These findings prompted the examination of CD4^+^ T cells in the tumor microenvironment (TME). Fifteen days post-treatment with LPS+PGPC+WTL, we observed a decrease in the amount of Foxp3^+^ CD4^+^ T cells in the TME, as compared to untreated mice **(Figure 1G, S1F).** Moreover, the ratio of IFNγ-producing to IL-10-producing CD4^+^ T cells in the TME increased >3-fold upon immunization with LPS+PGPC+WTL **(Figure 1G).** Thus, in elderly mice, CD4^+^ T cells are more important than CD8^+^ T cells for anti-tumor immunity, and the ability of CD4^+^ T cells to eradicate tumors correlates with a shift in the inflammatory status of these T cells in the TME.

### Inflammasomes, IL-1β and DC migratory activity are critical determinants of anti-tumor immunity in elderly mice

To determine if anti-tumor responses in elderly mice are dependent on endogenous DCs, as would be expected, we eliminated DCs from 68 weeks old mice. DC elimination was accomplished by reconstituting irradiated CD45.1 mice with bone marrow cells isolated from mice encoding the Diptheria Toxin Receptor (DTR) under control of the promoter for the gene *Zbtb46* (*Zbtb46*^DTR^). *Zbtb46* is commonly used to target transgenes for expression in DCs specifically. Six weeks after marrow transplant, chimerism was above 90% **(Figure S1G)** and conventional DCs were comprehensively eliminated following diphtheria toxin (DTx) injection **(Figure S1G)**.

In contrast, DTx injections had no impact on macrophage abundance in chimeric mice **(Figure S1G)**. We found that the tumor protection provided by LPS+PGPC was lost in immunized *Zbtb46*^DTR^ elderly mice **(Figure 1SH)**. This finding indicates that the mechanism by which hyperactivating stimuli induce tumor rejection from elderly mice is dependent on DCs. We therefore examined mechanisms intrinsic to DCs that impact protective immunity.

As IL-1β was necessary for protective immunity induced by hyperactivating stimuli **(Figure 1E-F, and Figure S1E)**, we performed genetic analysis of the upstream regulators of IL-1β production (and cell migration). We used 68 week old mice lacking NLRP3 (*Nlrp3^-/-^*) or mice lacking caspase-1 and caspase-11 (*Casp1/11^-/-^*). Both of these strains lack the ability to produce bioactive IL-1β^30, 40^. Similar analyses were performed on mice lacking CCR7 (*Ccr7^-/-^*), in which DCs display defects in migration to the skin draining lymph node (dLN). Mice were implanted with B16OVA tumors at 68 weeks of age. When tumors reached 2-3mm in size, mice were either left untreated or they were treated s.c. with LPS+PGPC+WTL. Wild type (WT) mice eradicated tumors after therapeutic immunization of LPS+PGPC+WTL. In contrast, LPS+PGPC+WTL treatments were ineffective when injected into *Nlrp3^-/-^, Ccr7^-/-^ or Casp1/11^-/-^*mice **(Figure 1H)**. These collective results indicate that DCs, inflammasomes, IL-1β and CCR7 are all necessary for the induction of anti-tumor immunity in elderly mice.

### DC hyperactivators induce TH1-skewed CD4^+^ T cell responses

The requirement of DCs and CD4^+^ T cells for anti-tumor responses in elderly mice can be explained if hyperactive DCs stimulate antigen-specific responses in this subset of T cells. However, our knowledge of how hyperactive DCs impact CD4^+^ T cell responses is minimal. To determine how distinct DC stimuli influence CD4^+^ T cell instruction *in vivo*, we first studied DCs in 8 week old mice. These mice were immunized s.c. with either PBS or with activating stimuli (LPS), hyperactivating stimuli (LPS+PGPC) or with pyroptotic stimuli (LPS+alum or alum alone). We monitored the behavior of endogenous DCs in the skin dLN 24 hours post-immunization. In contrast to mice immunized with PBS or LPS, the absolute number of DCs in the dLN increased strongly when mice were immunized with the hyperactivating stimulus LPS+PGPC **(Figure 2A).** Immunizations with the pyroptotic stimulus LPS+alum led to the lowest number of DCs in the dLN, a finding that is consistent with the idea that pyroptotic DCs lose the ability to migrate **(Figure 2A).** To assess functionality of the dLN DCs, we sorted these cells as CD11c^+^ MHC-II^+^CD11b^-^ live cells 24 hours following s.c. injection and cultured the cells for 24 hours on agonistic anti-CD40 coated plates. We then measured the secretion of the TH1 and TH2 driver cytokines, IL-12p70 and IL-10. dLN DCs from mice immunized with LPS+PGPC produced the highest amounts of IL-12p70, as compared to all other DC stimuli examined **(Figure 2B)**. In contrast to the ability to enhance IL-12p70, hyperactivating stimuli did not elicit the production of IL-10 **(Figure 2C)**. IL-10 production was most abundantly produced by DCs that were isolated from mice immunized with the pyroptosis-inducing stimuli LPS+alum or alum alone **(Figure 2C).** The levels of IL-12p70 secretion correlated with CD40 abundance on the surface of dLN DCs **(Figure 2D)**. The differences in IL-12p70 and IL-10 observed between PGPC-based and alum-based stimulations were notable, as both chemicals agonize NLRP3 inflammasomes. These findings illustrate that not all inflammasome stimuli are equivalent, and that stimuli that promote DC hyperactivation (PGPC) are more effective than those that promote pyroptosis (alum) in inducing signals important to TH1 cell mediated immunity.

**Figure 2.**
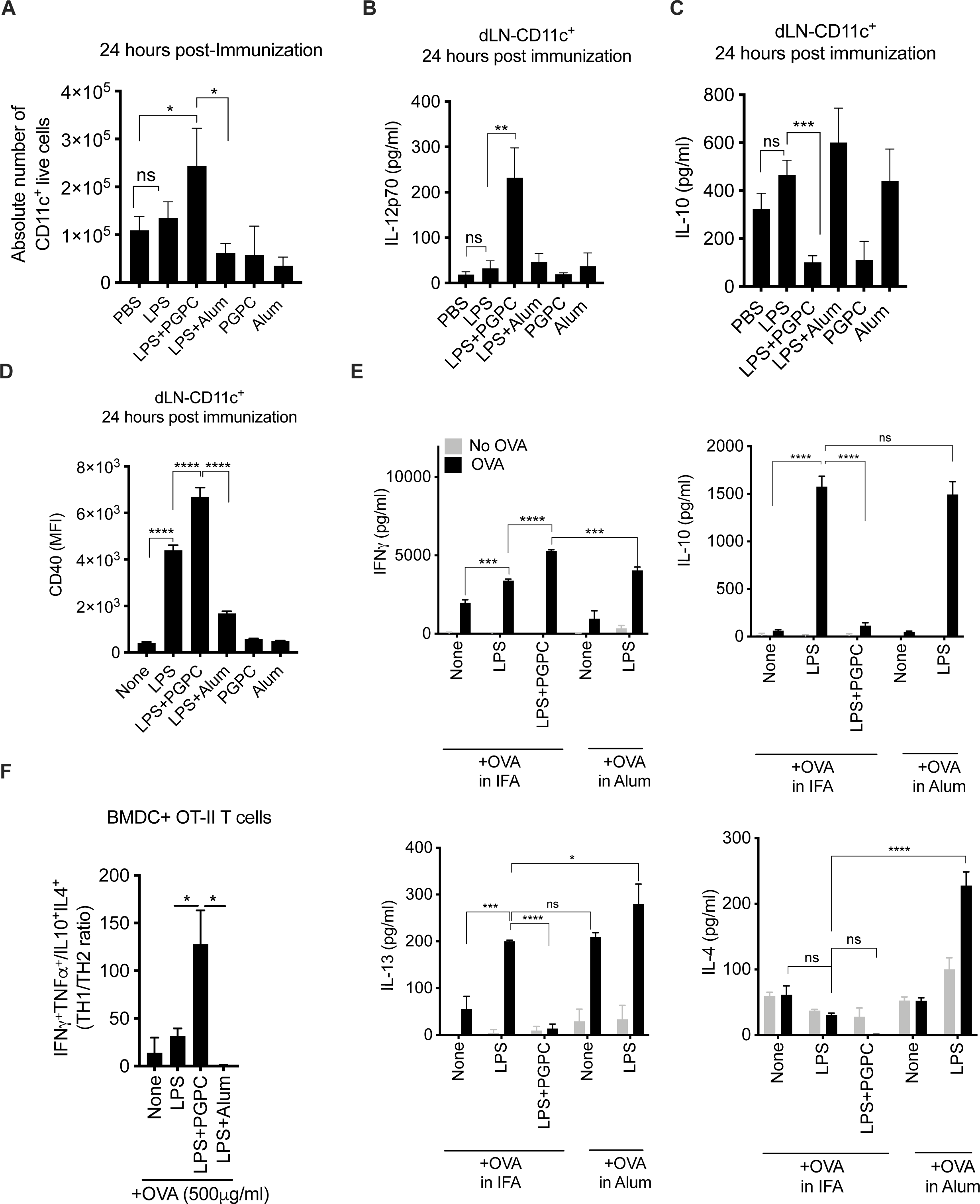
Hyperactive DCs drive TH1-skewed immune responses, with no evidence of TH2 immunity. (A-C) WT mice were injected subcutaneously (s.c.) on the right flank with PBS or LPS or PGPC or alum alone, or with LPS plus PGPC. Alternatively, mice were injected s.c. with Alum alone or LPS plus Alum. 24 hours post immunization, the skin draining lymph nodes (dLN) were isolated. (A) The absolute number of CD11c^+^ live cells was measured by flow cytometry. (B) CD11c^+^ cells were isolated as CD11c^+^MHC^-^II+CD11b^-^ then cultured onto a plate coated with agonistic anti-CD40 antibody for 24 hours. (B) IL-12p70 and (C) IL-10 secretion from dLN DCs were measured by ELISA. Means and SD of a triplicate is shown, and data are representative of at least 3 independent experiments. (D) CD40 expression by dLN DCs was measured by flow cytometry. Means and SDs of five mice are shown and are representative of 3 independent experiments. (E) WT mice were injected s.c. on the right flank with endofit-OVA protein either alone or with LPS, or with LPS plus PGPC emulsified in either incomplete Freud’s adjuvant (IFA) or in Alum as indicated. 40 days post immunization, the dLN were isolated. CD4^+^ were sorted then co-cultured with BMDC loaded (or not) with OVA. IFNγ, IL-10, IL-4 secretion was measured by ELISA. Means and SDs of four mice are shown and are representative of 3 independent experiments. (F) Bone marrow derived DCs (BMDCs) pretreated with indicated stimuli were loaded with OVA protein for 1h, then incubated for 4 days with OT-II naïve CD4+ T cells. 4 days later, T cells were stimulated for 5h with PMA plus ionomycin in the presence of brefeldin-A and monensin. The frequency of TH1 cells as TNFa^+^ IFNg^+^, and TH2 cells as IL-4^+^IL-10^+^ were measured by intracellular staining. Data are represented as the ratio of TH1/TH2 cells. Means and SDs of five mice are shown.

The enhanced ability of hyperactivating stimuli to upregulate factors important for TH1 differentiation prompted us to examine antigen-specific CD4^+^ T cell responses *in vivo*. Mice were immunized with the OVA antigen, either alone or with LPS, LPS+PGPC, LPS+alum, or alum alone. Forty days post-immunization, CD4^+^ T cells were isolated from the spleens of mice and re-stimulated *ex vivo* with naïve bone marrow derived DCs (BMDCs) that were loaded (or not) with OVA. Immunizations with LPS+PGPC led to the greatest amount of IFNγ production by responding CD4^+^ T cells, as compared to immunizations with LPS or pyroptotic stimuli (LPS+alum) **(Figure 2E).** Interestingly, the enhanced production of IFNγ did not correlate with enhanced production of TH2-associated cytokines. We observed no IL-10, IL-4 or IL-13 production by CD4^+^ T cells from mice immunized with LPS+PGPC **(Figure 2E)**. Thus, DC hyperactivators result in TH1-skewed T cell responses from endogenous T cells *in vivo* with minimal evidence of TH2 responses. In contrast to our findings with hyperactivating stimuli, LPS or LPS+alum immunizations led to a mixed TH1 and TH2 phenotype, as evidenced by the co-detection of IFNγ, IL-10, IL-4 and IL-13 after T cell restimulation by exposure to OVA-loaded GMCSF-BMDCs *ex vivo* **(Figure 2E).** Immunizations with OVA and alum alone led to a highly skewed IL-13 response, with no IFNγ being detected by responding T cells **(Figure 2E)**. These latter findings with alum corroborate studies showing that alum is a poor inducer of type I immunity^27, 41^, and that alum leads to TH2-skewed immune responses^28, 29, 42, 43^. In summary, these data demonstrate that stimuli that hyperactivate DCs selectively promote TH1 immune responses, with low TH2-induced immunity.

To examine the sufficiency of differentially stimulated DCs to drive distinct T cell activation programs, an *in vitro* co-culture system was utilized. We used BMDCs generated with (FMS-like tyrosine kinase 3 ligand) FLT3L, which represent conventional Type I (cDC1) and Type 2 (cDC2) cells. FLT3L-BMDCs were treated with the aforementioned stimuli. 24 hours post treatment, DCs were loaded (or not) with OVA and then exposed to naïve OT-II T cells. OT-II cells express a T cell Receptor (TCR) specific for an MHC-II restricted OVA peptide (OVA 323-339)^44^. The activities of the responding T cells were assessed by ELISA to determine T cell polarization towards TH1 responses (IFNγ, TNFα production) or TH2 responses (IL-10, IL-13 production). Regardless of DC activation state, OVA-treated DCs stimulated the production of IFNγ and TNFα from OT-II T cells **(Figure S2A)**. These results indicate that *in vitro*, TH1 responses are commonly induced, regardless of the activation state of the antigen presenting cells (APC). In contrast, TH2 responses were strikingly different when comparing DC activation states. Stimuli that induce DC activation (LPS) or pyroptosis (LPS+alum) promoted the release of large amounts of IL-10 and IL-13, whereas hyperactivating stimuli (LPS+PGPC) led to minimal production of these TH2-associated cytokines **(Figure S2A)**.

Intracellular staining of single cells for TH1 (IFNγ and TNFα) and TH2 (IL-4 and IL-10) cytokines permitted the identification of TH1 and TH2 cells generated by different DC stimuli. This analysis revealed that hyperactive DCs induced a strong skewing of individual T cells towards the IFNγ-expressing TH1 lineage **(Figure 2F and S2B)**. The ratio of TH1 (IFNγ^+^ TNFα^+^) to TH2 (IL-10^+^ IL-4^+^) cells stimulated under hyperactivating conditions was greater than 100:1 **(Figure 2F and S2B)**. In contrast, all other DC stimuli induced a mixed T cell response, with pyroptotic stimuli leading to a nearly 1:1 ratio of TH1 to TH2 cells **(Figure 2F and S2B)**. In addition, intracellular staining of the TH2-lineage transcription factor GATA3 along with IL-4 indicated that while pyroptotic stimuli induce the highest percentage of GATA3^+^ IL-4^+^ cells, hyperactivating stimuli induce 4-fold less TH2 cells **(Figure S2C)**.

We determined if the ability of hyperactive DCs to induce TH1 skewed responses extended to the functions of antigen experienced (*i.e.* memory) T cells. We isolated CD4^+^ CD44^+^ memory T cells from mice that were previously immunized with OVA in Incomplete Freund’s Adjuvant (IFA). T cells were then cultured with OVA-loaded FLT3L-BMDCs that were stimulated as described above. As compared to other DC stimuli, LPS+PGPC stimulated DCs induced the highest amount of IFNγ expression among Tbet^+^ T cells **(Figure 3A)**. Collectively, these *in vitro* and *in vivo* datasets support the conclusion that distinct DC stimuli elicit unique T cell activation states, with hyperactivating stimuli being the most potent at inducing TH1 cell activities and restimulating antigen-experienced (*i.e.* memory) TH1 cells.

**Figure 3.**
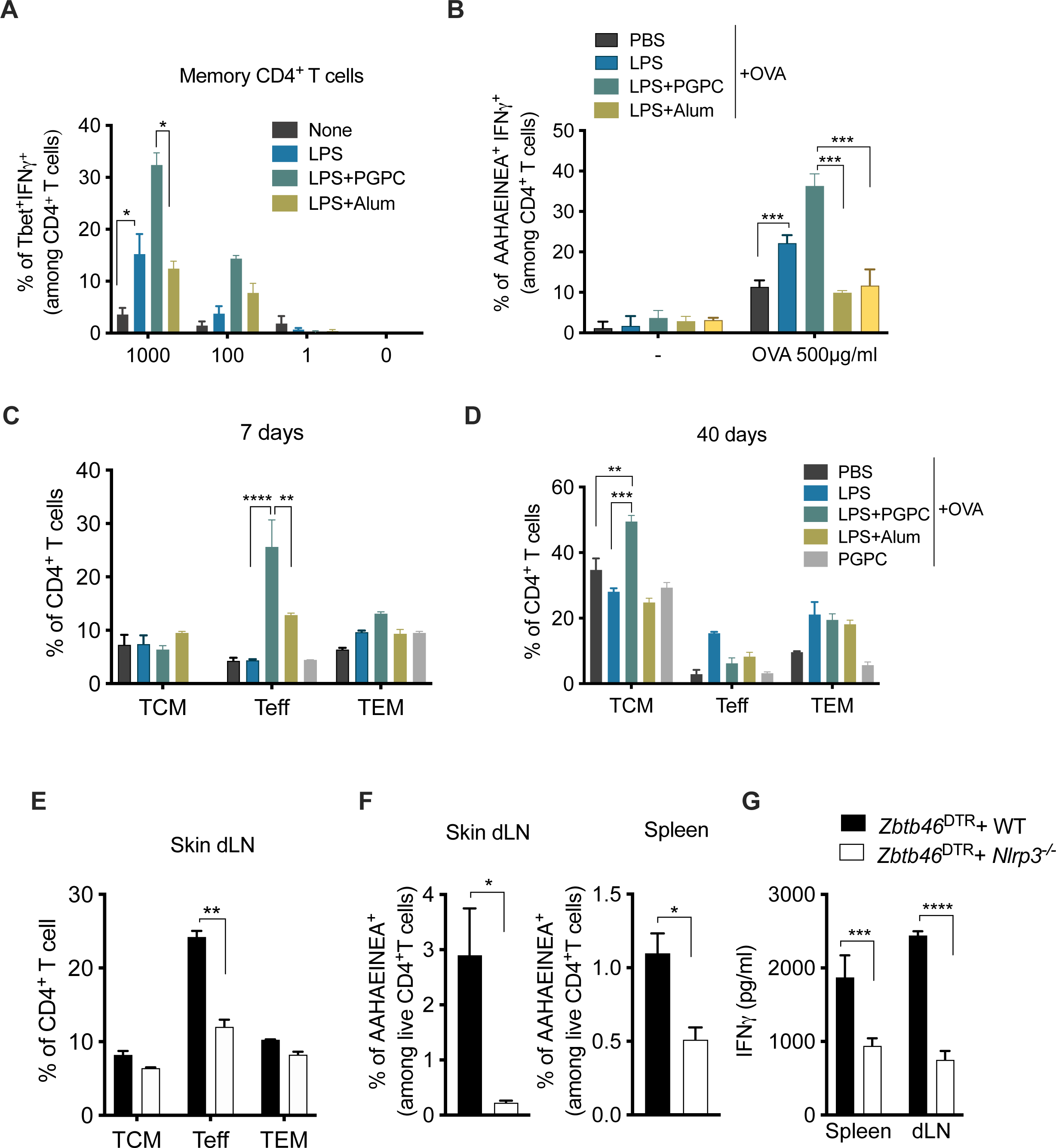
DC hyperactivators enhance CD4^+^ reactivation of already committed TH1 cells and accelerate memory T generation. (A) Mice received three boost injections s.c. with OVA emulsified in IFA every 7 days. 7 days after the last boost, splenic memory CD4^+^ T cells were co-cultured with DCs that were pretreated with indicated stimuli and loaded or not with as serial dilution of OVA. 5 days post co-culture, the percentage of Tbet^+^IFNg^+^T cells was measured by flow cytometry (B) Mice were injected s.c. on the right flank with OVA either alone or with LPS, or with LPS plus PGPC all emulsified in IFA, or with LPS plus Alum. Some mice were injected with LPS+PGPC without OVA antigen (no OVA). 7 days post-immunization, CD4^+^ T cells were sorted from the skin draining lymph nodes (dLN) of immunized mice, and co-cultured with BMDCs loaded (or not) with OVA for 5 days at a ratio of 1:10 (DC: T cell). The percentage of AAHAEINEA^+^ IFNγ+ among CD4^+^live T cells was measured using OVA peptide tetramer staining followed by an intracellular IFNγ staining. Means and SDs of five mice are shown. (C-D) WT mice were injected s.c. on the right flank with endofit-OVA protein either alone or with LPS or with PGPC, or with LPS plus PGPC all emulsified in incomplete Freud’s adjuvant (IFA). Alternatively, mice were injected s.c. with Alum alone or LPS plus Alum. (C) Seven days and (D) forty days post immunization, skin dLN were dissected and T cells were enriched using anti-CD4 magnetic beads. The percentage of T effector cells (Teff) as CD44^lo^CD62L^lo^, T effector memory cells (TEM) as CD44^hi^CD62L^-^, and T central memory cells (TCM) as CD44^hi^ CD62L^hi^ are represented by gating on CD3^+^CD4^+^ live cells. (E-G) CD45.1 mice were irradiated then reconstituted with mixed Bone marrow of the genotypes indicated. Six weeks post-reconstitution, chimera mice were injected with diphteria toxin (DTx) 3 times a week for a total of 9 DTx injections. Chimeric mice were then immunized s.c. on the right flank with OVA with LPS plus PGPC. (E) Seven days post-immunization with OVA with LPS and PGPC, the percentage of Teff, TEM, TCM, and T naive cells in the skin dLN was measured by flow cytometry. (F) The percentage of AAHAEINEA^+^ among CD4^+^ live T cells from the spleen and dLN was measured using OVA peptide tetramer staining. Means and SDs of five mice are shown. (G) CD4^+^ T cells were sorted from the skin dLN of immunized mice then co-cultured with BMDC loaded with OVA at a ratio of 1:10 (DC: T cell) for 4 days. IFNγ, IL-10 and IL-4 release in the supernatants of BMDC-CD4^+^ T cells co-culture were measured by ELISA. Means and SDs of five mice are shown.

### DC hyperactivators accelerate memory CD4^+^ TH1 cell differentiation in an NLRP3-dependent manner

Based on the ability of hyperactivating stimuli to induce TH1-skewed T cell responses, we determined if we could identify individual endogenous OVA-specific T cell responses in mice. Mice were injected with OVA alone or with LPS, LPS+alum, or LPS+PGPC. Alternatively, mice were immunized with LPS+PGPC without the OVA antigen. 7 days post immunization, CD4^+^ T cells were isolated from the skin dLN and were re-stimulated *ex vivo* with naïve BMDCs loaded (or not) with OVA to enrich the OVA-specific T cell subset. T cell effector function of OVA-specific TH1 cells was assessed by intracellular staining for IFNγ, and TCR specificity was assessed by staining with I-A(d) OVA peptide (OVA 329-337) tetramers. We found that PGPC-based immunizations generated the highest frequency of tetramer^+^ IFNγ^+^ responses upon CD4^+^ T cell re-stimulation with OVA antigen **(Figure 3B)**. Pyroptotic stimuli (LPS+alum) were the weakest inducers of antigen-specific IFNγ responses (**Figure 3B)**. These data reveal hyperactivating stimuli as effective inducers of endogenous OVA-specific TH1 T cell responses in mice.

The enhanced TH1 activities associated with hyperactivating stimuli could be explained by an altered T cell differentiation trajectory after immunization. To examine this possibility, we monitored memory T cell generation and effector memory T cell responses *in vivo*. Mice were immunized with OVA alone and the aforementioned DC stimuli. 7- and 40-days post-immunization, memory and effector T cell generation in the dLN was assessed by flow cytometry using CD44 and CD62L markers that distinguish T effector cells (Teff) as CD44^low^CD62L^low^, T effector memory cells (TEM) as CD44^hi^CD62L^low^, and T central memory cells (TCM) as CD44^hi^CD62L^hi^ ^45, 46^. Seven days post-immunization, hyperactivating stimuli were superior to activating stimuli at inducing CD4^+^ Teff cells **(Figure 3C)**. At this early time point, both stimuli induced low but comparable amounts of CD4^+^ TEM cells **(Figure 3C)**. Forty days post-immunization, ample TCM cells were observed in mice exposed to hyperactivating stimuli, whereas these cells were less abundant in mice immunized with OVA alone or with LPS **(Figure 3D)**. These data indicate that the hyperactivating stimulus PGPC accelerate the magnitude of effector and memory T cell generation.

To define mechanisms by which hyperactivating stimuli accelerate CD4^+^ T cell differentiation, we identified DC-intrinsic determinants of this process. Previous studies showed that antigen-specific T cell responses can be enhanced by co-immunization with recombinant IL-1β^47^, a cytokine whose bioactivity is naturally controlled by inflammasomes. However, the role of inflammasomes within DCs for CD4^+^ T cell activities is poorly defined. To determine the role of inflammasomes within endogenous DCs in mediating hyperactivation-based T cell activities, we generated mixed chimeras in mice as described^30, 48^. Briefly, CD45.1 mice were irradiated and reconstituted with mixed bone marrow (BM) using 80% BM cells isolated from *Zbtb46*^DTR^ mice mixed with 20% BM cells isolated from WT, or *Nlrp3*^-/-^ mice on a CD45.2 background. Six weeks post-reconstitution, the efficacy of reconstitution was above 92% efficiency. Reconstituted mice were then injected i.p. with DTx to deplete Zbtb46^+^ cDCs, giving rise to mice that harbor either WT or inflammasome-deficient (*Nlrp3^-/-^*) DCs. The resulting mice were then s.c. immunized with OVA plus LPS+PGPC. Seven days later, CD4^+^ T cell responses from the skin dLN and spleen were assessed. We found that the abundance of Teff CD4^+^ T cells in the dLN was reduced in mice harboring DCs that cannot become hyperactive (*Nlrp3^-/-^* chimera mice), as compared to mice harboring WT DCs **(Figure 3E, Figure S3A)**. To monitor antigen specificity, we measured CD4^+^ T cells in WT and *Nlrp3^-/-^* chimera mice using the OVA peptide (AAHEINEA) tetramers. We identified lower AAHEINEA^+^ T cells in the skin dLN and spleen of immunized *Nlrp3*^-/-^ chimera mice, as compared to WT chimeras **(Figure 3F, Figure S3B)**. Moreover, the frequency of antigen-specific T cells observed 7 days post-immunization correlated with enhanced IFNγ responses of CD4^+^ T cells **(Figure 3G)**. Collectively, these data demonstrate the importance of NLRP3 within hyperactive DCs to accelerate the differentiation and functions of antigen-specific CD4^+^ T cells.

### DC hyperactivators induce CD4^+^ TH1 cells that persist into old age and eliminate tumors

A hallmark of adaptive immunity is the ability to provide long-term protection from a previously encountered threat to the host. Thus, we determined the longevity of the T cell responses induced by distinct DC activation states. Eight week old mice were injected s.c. with OVA alone or with LPS, LPS+alum or the DC hyperactivator LPS+PGPC. Alternatively, *Nlrp3*^-/-^ mice were immunized s.c. with OVA and LPS+PGPC. One year later, at 60 weeks post immunization, all mice received a s.c. boost with OVA alone emulsified in IFA. One week post-OVA re-exposure, all mice were implanted with B16OVA cells. Mice immunized with OVA alone or OVA+LPS succumbed to tumor growth **(Figure 4A)**. In contrast, immunizations with LPS+PGPC induced a rejection of B16OVA tumors in 80% of immunized mice. This protection was dependent on NLRP3, as *Nlrp3*^-/-^ mice were unable to survive the tumor challenge **(Figure 4A)**. LPS+alum immunizations with OVA were also unable to mediate tumor rejection, indicating that pyroptosis-inducing stimuli are not as effective as DC hyperactivators at inducing long-lived anti-tumor immunity **(Figure 4A)**.

**Figure 4.**
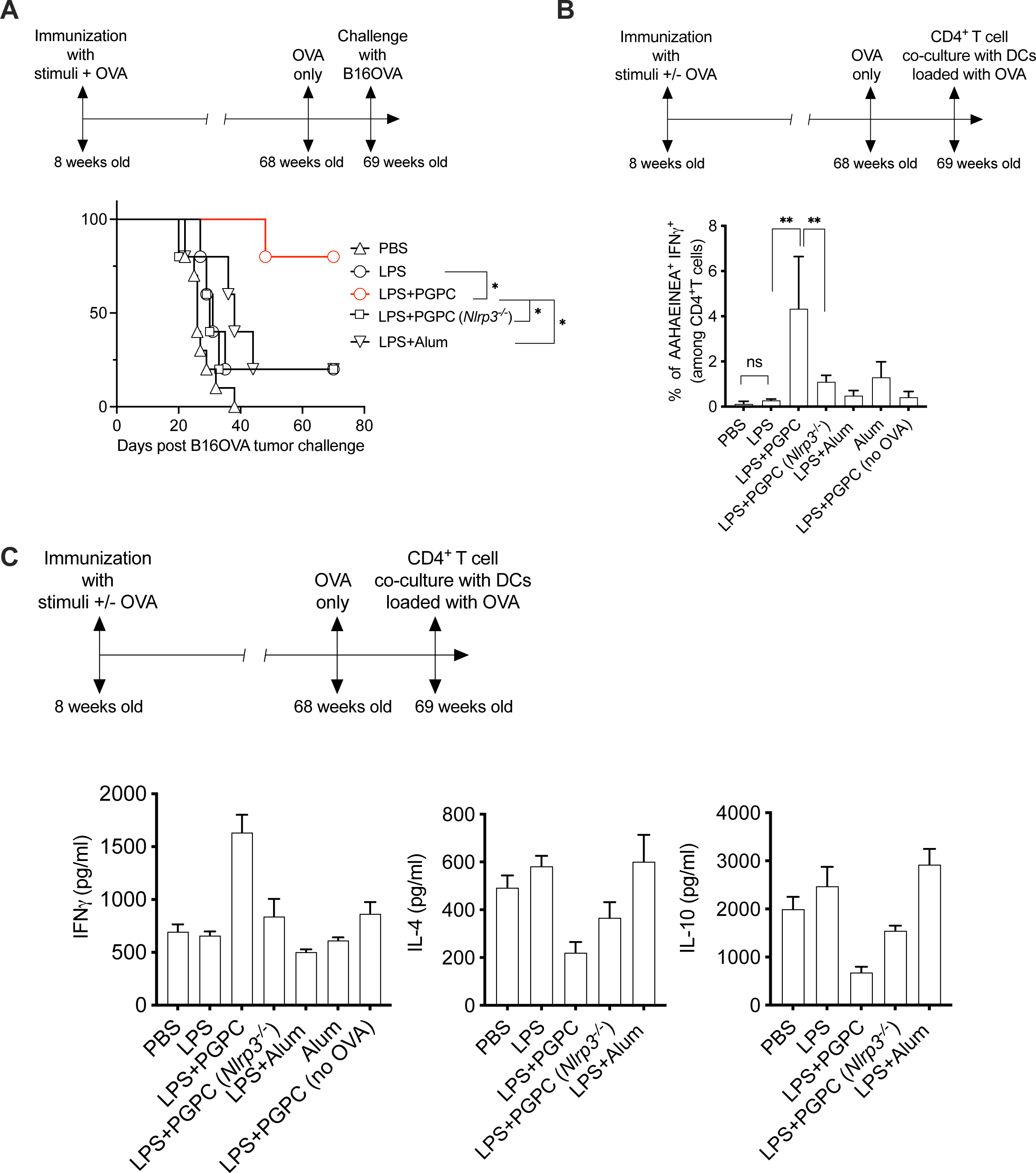
Hyperactive stimuli induce long-lived antigen specific CD4^+^ T cells that eradicate tumors during aging. (A-C) 8 weeks old mice were injected s.c. on the right flank with OVA either alone or with LPS, or with LPS plus PGPC all emulsified in IFA, or with LPS plus Alum. Alternatively, *Nlrp3*^-/-^mice were injected with OVA with LPS plus PGPC emulsified in IFA. 60 weeks later, mice were rechallenged s.c. on the right flank with OVA alone emulsified in IFA. In B, some mice were injected with LPS+PGPC without OVA antigen (no OVA). (A) One week post OVA rechallenge, mice were injected with B16OVA cells on the left flank. Percentage of mice survival was monitored every 2 days (n=5 mice per group). (B-C) 60 weeks later, mice were rechallenged s.c. on the right flank with OVA alone emulsified in IFA. CD4^+^ T cells were isolated one week after OVA re-exposure, and co-cultured with BMDCs loaded with OVA for 4 days. (B) The percentage of AAHAEINEA^+^ IFNγ^+^ among CD4^+^ live T cells was measured using OVA peptide tetramer staining followed by an intracellular IFNγ staining. (C) IFNγ, IL-4 and IL-10 release in the supernatant were measured by ELISA. Means and SDs of five mice are shown.

Due to the central importance of CD4^+^ T cells in protective immunity in elderly mice (Figure 1), we examined antigen specificity of CD4^+^ T cells within mice over a year after immunization. Eight weeks old mice were immunized with stimuli as mentioned above in the presence or absence of OVA. Mice then received a boost injection with OVA alone emulsified in IFA 60 weeks later. One week after the boost, these 69 week old mice were sacrificed and CD4^+^ T cells were sorted from the lymph node that drains the s.c. site of boost immunization. Sorted T cells were then re-activated *in vitro* by BMDCs loaded with OVA. Tetramer staining revealed that mice immunized with LPS+PGPC+OVA displayed the highest amount of OVA-specific CD4^+^ T cells in the dLN 60+ weeks post immunization, as compared to mice immunized with OVA alone or with LPS or with LPS+alum **(Figure 4B**). This ability of DC hyperactivators to maintain long-lived T cells was dependent on inflammasomes, as LPS+PGPC+OVA immunizations were ineffective when injected into NLRP3-deficient mice **(Figure 4B)**. To determine if the T cell responses were derived from the initial immunizations (as compared to primary T cell responses elicited by the boost), we examined mice that received a primary immunization with LPS+PGPC in the absence of OVA (but received an OVA injection 60 weeks post-immunization). Under these conditions, minimal OVA-specific CD4^+^ T cells were detected upon *in vitro* restimulation with OVA-loaded BMDCs **(Figure 4B)**. These data indicate that NLRP3 inflammasomes are required for DCs to induce antigen-specific T cells that persist for over a year.

Prior studies have found that CD4^+^ T cells isolated from elderly mice are promiscuous producers of IFNγ^49^. This process is a hallmark of inflammaging^31^. Consistent with this idea, we found that, regardless of the immunization stimulus used at 8 weeks of age, all CD4^+^ T cells examined from elderly mice secreted a basal amount of IFNγ **(Figure 4C)**. Despite this basal IFNγ producing activity of T cells from elderly mice, we found that immunizations early in life with hyperactivating stimuli induced the highest amounts of IFNγ secretion upon co-culture with OVA-loaded BMDCs, but the lowest amounts of IL-4 **(Figure 4C),** from T cells isolated from elderly mice. The ability of LPS+alum to induce high amounts of TH2 CD4+ T cells also persists into the later stages of life **(Figure 4C)**.

Genetic analysis revealed that early life (8 week) immunizations of *Nlrp3^-/-^* mice with LPS+PGPC were ineffective in inducing antigen-specific T cell responses that persisted into old age **(Figure 4B)**. In addition, elderly *Nlrp3^-/-^*mice were unable to induce TH1-skewed T cell responses after an early life immunization with LPS+PGPC+OVA **(Figure 4C)**. Collectively, these data suggest that NLRP3 inflammasomes are necessary for DC hyperactivators to induce memory TH1 skewed responses that persist into old age.

### Hyperactivating stimuli correct immune activities that are defective in elderly DCs

The ability to successfully immunize individuals later in life has been a challenge, as the aging immune system is considered defective in several areas^31^. The ability of DC hyperactivators to confer anti-tumor immunity upon immunizations of elderly mice (Figure 1) suggested that age-associated defects in DC functions may be corrected by DC hyperactivators. To test this idea, we examined properties of DCs that were prepared from young and elderly mice. This analysis revealed several categories of age-dependent DC activities.

The first category represented DC activities that operate similarly in young and elderly mice. We found that any stimulation performed in the presence of LPS, regardless of age, led to compared amounts of TNFα production by FLT3L-BMDCs **(Figure S4A)**. Similarly, IL-1β production was age-independent, and was only observed when cells were stimulated with NLRP3 inflammasome agonists such as LPS+PGPC or LPS+alum **(Figure S4B)**. The pyroptosis indicator lactate dehydrogenase (LDH) was only released into the extracellular space upon LPS+alum treatment, regardless of age **(Figure S4C)**. Thus, inflammasome-dependent responses and TLR-induced TNFα production represent age-independent responses.

The second category represented DC activities that were defective in old mice and not corrected by DC hyperactivators. For example, all LPS-containing stimuli induced greater amounts of IL-10 production (to varying extents) from young DCs, as compared to elderly DCs. This defect in IL-10 production was not corrected by any DC stimulus and IL-10 production was nearly absent from cells stimulated with LPS+PGPC **(Figure 5A)**.

**Figure 5.**
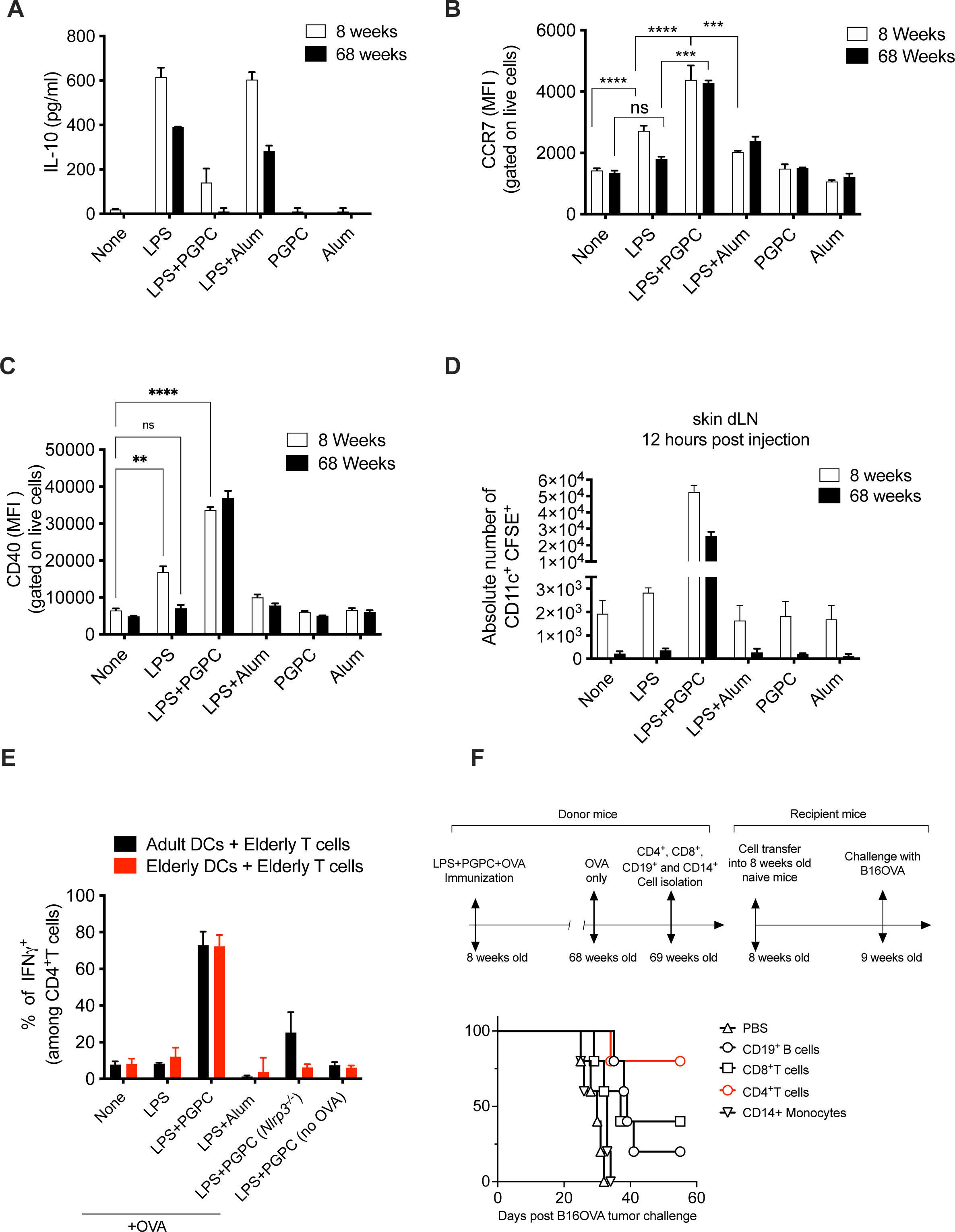
Hyperactive stimuli restore CD4^+^T cell stimulatory activities in aging DCs and induce tumor rejection in aging mice. (A-B) 8 weeks or 68 weeks old FLT3L-derived BMDCs were either left untreated (‘‘None’’) or treated with LPS alone, or Alum alone, or PGPC alone for 24h, or BMDCs were primed for 3h with LPS, then treated with indicated stimuli for 21h. (A) IL-10 release was monitored by ELISA. (B-C) The mean fluorescence intensity (MFI) of surface (B) CCR7 and (C) CD40 (among CD11c^+^live cells) was measured by flow cytometry. (D) DCs were labeled with CFSE and injected subcutaneously (s.c.) on the right flank in CD45.1 mice. 12 hours post injection, the skin draining lymph nodes (dLN) were isolated. The absolute number of CD45.2^+^CFSE^+^ among CD11c^+^ live cells was calculated by flow cytometry. Means and SDs from five mice are shown, and data are representative of at least three independent experiments. (E) 8 weeks or 68 weeks old BMDCs were generated from WT mice and treated as in A. Alternatively, BMDCs were generated from *Nlrp3^-/-^* mice. Treated DCs were loaded or not (NO OVA) with OVA protein for 1 hour then co-cultured with CD4^+^ T cells that were isolated from the 68 weeks old OVA immunized mice. 4 days post co-culture, IFNγ expression in CD4^+^T cells co-culture was measured by intracellular staining. Means and SDs of five mice are shown. (F) 8 weeks old mice were injected s.c. on the right flank with OVA in combination with LPS plus PGPC all emulsified in IFA. 60 weeks later, mice were rechallenged s.c. on the right flank with OVA alone emulsified in IFA. CD4^+^, CD8^+^ T cells, CD19^+^ B cells and CD14^+^ monocytes were isolated from aging mice that were immunized with OVA and LPS plus PGPC. Isolated cells were activated for 24 hours before their injection intravenously into 8 weeks old naïve mice. One week post cell transfer, all mice were challenged with B16OVA cells on the left flank. Percentage of mice survival was monitored every 2 days (n=5 mice per group).

The third category represented DC activities that were defective in old mice but were corrected by treatments with DC hyperactivators. For example, the ability of LPS to induce CCR7 **(Figure 5B)** and CD40 **(Figure 5C)** expression diminished with age, as was the ability of LPS to induce IL-12p70 production following DC culture on plates coated with agonistic anti-CD40 antibodies **(Figure S4D)**. All three of these defects were corrected when DCs were treated with LPS+PGPC, but not with other DC stimuli. The correction of the age-associated defects in CCR7 expression raised the question of DC migratory activity. To test this, young and elderly DCs were treated as above, then stained with carboxyfluorescein succinimidyl ester (CFSE) and injected s.c. into mice on a CD45.1 background, as described^30^. Twenty hours post-injection, we measured the absolute number of DCs that migrated to the dLN as CD11c^+^ CFSE^+^ among CD45.2^+^ living cells. We found that untreated or LPS-treated elderly DCs were defective in migration, as compared to similarly stimulated DCs from young mice **(Figure 5D).** This age-associated migration defect has been reported in mice and humans^2, 50, 51^. When young or elderly DCs were treated with the LPS+PGPC prior to s.c. injection, migration to the dLN was enhanced substantially and far exceeded the migratory activity of any other stimulus examined **(Figure 5D)**. While we observed that elderly hyperactive DCs were less migratory than young hyperactive DCs (both of which were treated with LPS+PGPC), the amount of migration observed by hyperactive DCs was 50 times greater than any migratory activity observed for DCs that were treated with LPS alone. The pyroptotic stimulus LPS+alum, in contrast, failed to induce any migration by young or elderly DCs. Overall, these data reveal a strategy to alleviate several age-associated defects in DC activities that contribute to efficient induction of adaptive immunity.

### Effective immunizations of elderly mice are directed by NLRP3 in hyperactive DCs

Based on the ability of DC hyperactivators to correct age-associated defects in DC functions, we examined the induction of T cell mediated adaptive immunity. We found that neither LPS nor LPS+alum treated DCs had the ability to induce OVA-specific IFNγ production from T cells that were isolated from elderly mice that were previously immunized with OVA in IFA. Only DCs treated with LPS+PGPC induced large amounts of IFNγ secretion from elderly T cells **(Figure 5E**). This IFNγ response was dependent on NLRP3 inflammasomes, as *Nlrp3^-/-^* DCs were unable to elicit IFNγ production **(Figure 5E**). When mice were immunized with LPS+PGPC in the absence of OVA, CD4^+^ T cells did not secrete IFNγ under any condition examined, indicating that antigen-specific T cell responses were observed **(Figure 5E)**. These data reveal the unique ability of hyperactive DCs to elicit T cell responses from elderly mice. The inability of LPS- or alum-based adjuvants to induce T cell responses in old mice may underlie problems associated with protective immunity in elderly humans.

To determine if the T cells generated by DC hyperactivators were sufficient to confer protective immunity, we transferred select immune cells into a heterologous host mouse. We immunized mice at 8 weeks of age with OVA and LPS+PGPC. At 69 weeks of age, we isolated CD4^+^ T cells, CD8^+^ T cells, CD19^+^ B cells and CD14^+^ monocytes from these mice. Each of these immune cell types was transferred into a fresh set of 8 weeks old mice. One-week after cell transfer, recipient mice were challenged with a lethal dose of B16OVA cells. While CD19^+^ and CD8^+^ cells induced a minor anti-tumor protection, CD4^+^ T cells transfer was sufficient to induce tumor rejection in 80% of recipient young mice **(Figure 5F)**. These data provide a mechanistic and functional explanation for how hyperactivating DC stimuli protect young and elderly mice from cancer, as this DC activation state is uniquely capable of eliciting long-lived memory CD4^+^ T cells.

### Hyperactivators correct age-associated DC defects and induce NLRP3- and Gasdermin D dependent TH1-skewed responses from young and old humans

All knowledge of the hyperactive DC state elicited by PGPC derives from studies of murine cells. Whether LPS+PGPC promotes a state of hyperactivation in human DCs is unknown. To test this, we generated primary human monocyte-derived DCs (moDCs) by culturing monocytes with GMCSF and IL-4. The purity of moDCs was defined as the percentage of CD14^neg^CD11c^+^CD209^+^, and was above 93% in all samples. moDCs were left untreated or treated with the DC stimuli used throughout this study. 24 hours post-stimulation, we tested moDCs for hallmarks of hyperactivation, such as their ability to secrete IL-1β from living cells. We found that only LPS+PGPC, LPS+alum and LPS+nigericin treatments induced the release of IL-1β **(Figure 6A upper panel)**. Alum and nigericin treatments, as expected, coupled IL-1β released with cell death, as revealed by LDH release **(Figure 6A lower panel)**. In contrast, LPS+PGPC induced IL-1β release in the absence of LDH release in all human subjects examined **(Figure 6A lower panel)**. The release of IL-1β was dependent on the concentration of PGPC **(Figure 6B)**.

**Figure 6.**
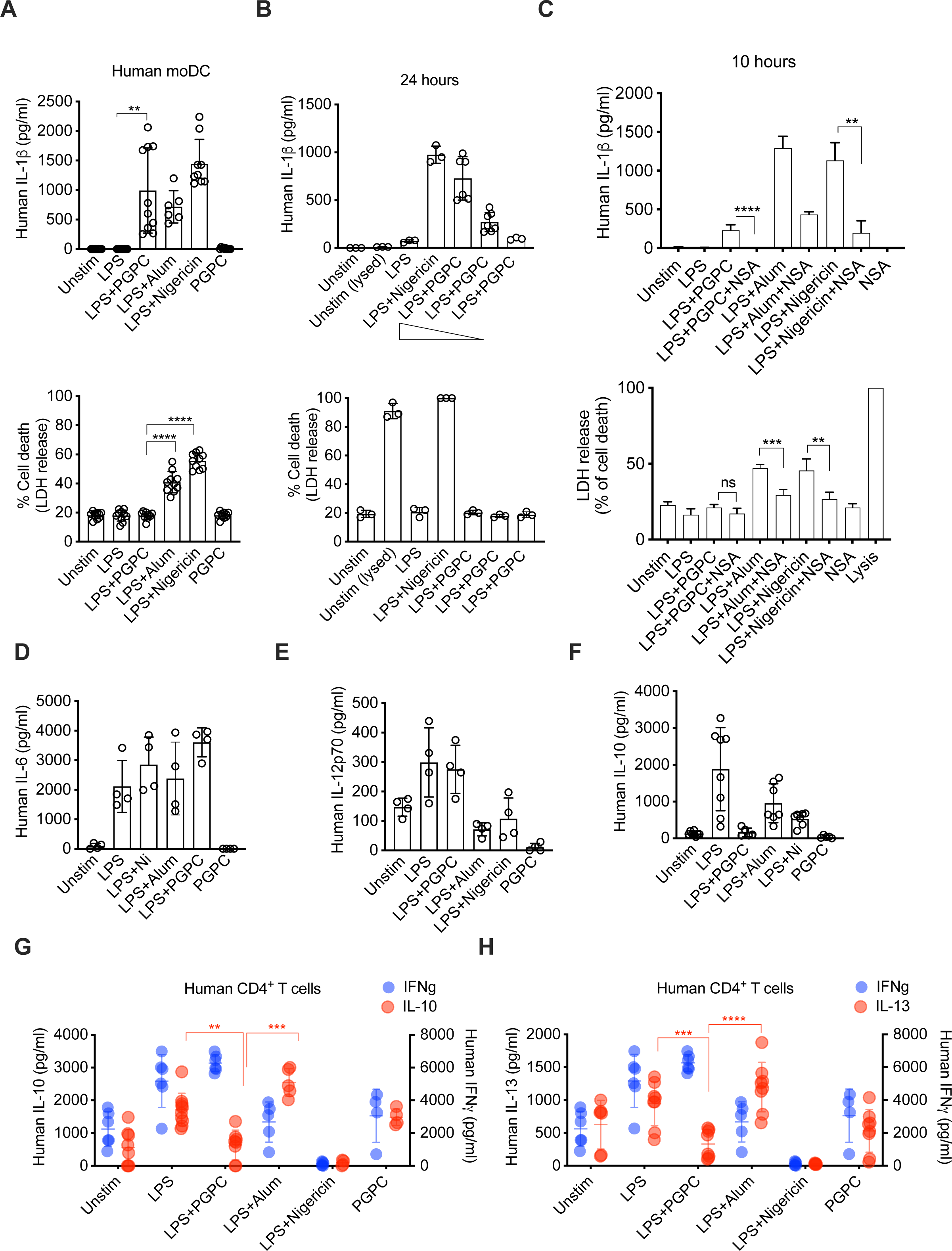
Human moDCs achieve a state of hyperactivation and drive TH1-skewed immune responses. (A-B) Human moDC generated from healthy adult donors were either left untreated (‘‘Unstim’’) or treated with LPS alone, or alum alone, or PGPC alone for 24h, or DCs were primed for 3h with LPS, then treated with indicated stimuli for 21h. (A) IL-1β in upper panel was measured by ELISA from 6-10 donors and LDH release in bottom panel is indicated. (B) Human moDCs were treated as above for 21h. For hyperactivation, DCs were primed with LPS for 3 hours then treated with PGPC at 100ug/ml or 50 ug/ml or 25 ug/ml for 21 hours. (upper panel) IL-1β and (bottom panel) LDH release was monitored by ELISA. Means and SDs of at least 3 donors is represented. (C) moDCs were either left untreated (‘‘Unstim’’) or treated with LPS alone, or alum or necrosulfonamide (NSA) alone for 10 hours. Alternatively, moDCs were primed for 4h with LPS then treated with PGPC or alum for 6h with or without NSA. (Upper panel) IL-1β, and (bottom panel) LDH and release was measured. Means and SDs of 4 donors is represented. (D) moDC were treated as above with indicated stimuli. (D) IL-6 (E) IL-10 and (F) IL-12p70 release was monitored by ELISA 24 hours post stimulation. Means and SDs of 4-8 donors is represented. (G-H) moDC were treated as above, then cultured with allogeneic T cells for 5 days. (G) Human IFNγ and IL-10, and (H) Human IFNγ and IL-13 secretion was measured by ELISA. Means and SDs of 4-8 donors is represented.

In murine cells, DC hyperactivators mediated IL-1β release via the actions of the inflammasome effector protein gasdermin-D (GSDMD), which forms plasma membrane pores that serve as conduits for IL-1β release^52^. Under pyroptosis-inducing conditions, GSDMD pores are formed in great excess to those formed under hyperactivating conditions, leading to plasma membrane rupture and cell death^52^. To assess whether IL-1β release from human moDCs occurs via GSDMD, we performed moDCs stimulations with or without the GSDMD chemical inhibitor necrosulfonamide (NSA)^53^. Under all conditions in which IL-1β and LDH release were observed, NSA treatments prevented these activities **(Figure 6C)**. These results suggest that the process by which mammalian cells achieve a hyperactive state is conserved in murine and human DCs, and that GSDMD is a mediator of these events in both species.

We assessed the impact of different DC stimuli on the ability of moDCs to produce cytokines including IL-6, IL-12p70 and IL-10. We found that all stimuli containing LPS induced similar amount of IL-6 secretion from DCs **(Figure 6D).** Similarly, stimulations with LPS and LPS+PGPC elicited compared amounts of IL-12p70 secretion from moDCs **(Figure 6E)**. However, the pyroptotic stimuli LPS+alum or LPS+nigericin did not induce moDCs to secrete IL-12p70 **(Figure 6E)**. Interestingly, while LPS treated moDCs produced IL-10, LPS+PGPC strongly diminished IL-10 secretion by moDCs **(Figure 6F)**. These data were similar to what we have observed in murine DCs **(Figure S4A)**. LPS+alum induced high levels of IL-10, whereas LPS+nigericin treatment was poor at inducing cytokine secretion from moDCs **(Figure 6F)**, likely due to moDC death **(Figure 6A)**. Thus, similar to murine cells, hyperactive moDCs display several activities that would predict T cell stimulatory capacity.

We assessed the impact of stimulated DCs on T cell functions using the mixed leukocyte reaction (MLR). Stimulated moDCs were cultured with allogeneic naïve T cells for 5 days and TH1 (IFNγ), and TH2 (IL-13 and IL-10) cytokine secretion was assessed. Unstimulated and LPS-treated moDCs promoted T cells to secrete IFNγ, IL-10 and IL-13 **(Figure 6G-H)**, indicating a mixed TH1-TH2 response. Hyperactive moDCs treated with LPS+PGPC produced similar amounts of IFNγ as LPS-treated moDCs (**Figure 6G)**, but secreted minimal IL-13 and IL-10 **(Figure 6G-H)**. This TH1-skewed phenotype induced by hyperactive DCs contrasted with the TH2-skewed response induced by the pyroptotic stimulus LPS+alum. LPS+alum elicited high amounts of IL-10 and IL-13 secretion, but low amounts of IFNγ **(Figure 6G-H)**. moDCs treated with LPS+nigericin were unable to stimulate T cells. Collectively, these data indicate that in contrast to other DC activation states, hyperactive DCs induce TH1 skewed responses while dampening TH2 responses.

The effectiveness of hyperactivating stimuli in human moDCs provided the mandate to explore their activities on aging moDCs. We generated moDCs from female or male individuals over 70 years of age and exposed these cells to DC stimuli. Assays for IL-6 and IL-1β production revealed that all LPS-containing stimuli elicited production of the former and all inflammasome stimuli elicited the latter **(Figure 7A)**. LPS+PGPC induced IL-1β production was sensitive to the NLRP3 inhibitor MCC950^54, 55^, indicating that IL-1β production was inflammasome-mediated **(Figure 7D)**. Pyroptotic stimuli (LPS+alum or LPS+nigericin) were able to couple IL-1β production with cell death, as assessed by LDH release, whereas LPS+PGPC induced comparable amounts of IL-1β as pyroptotic cells from living moDCs **(Figure 7C)**.

**Figure 7.**
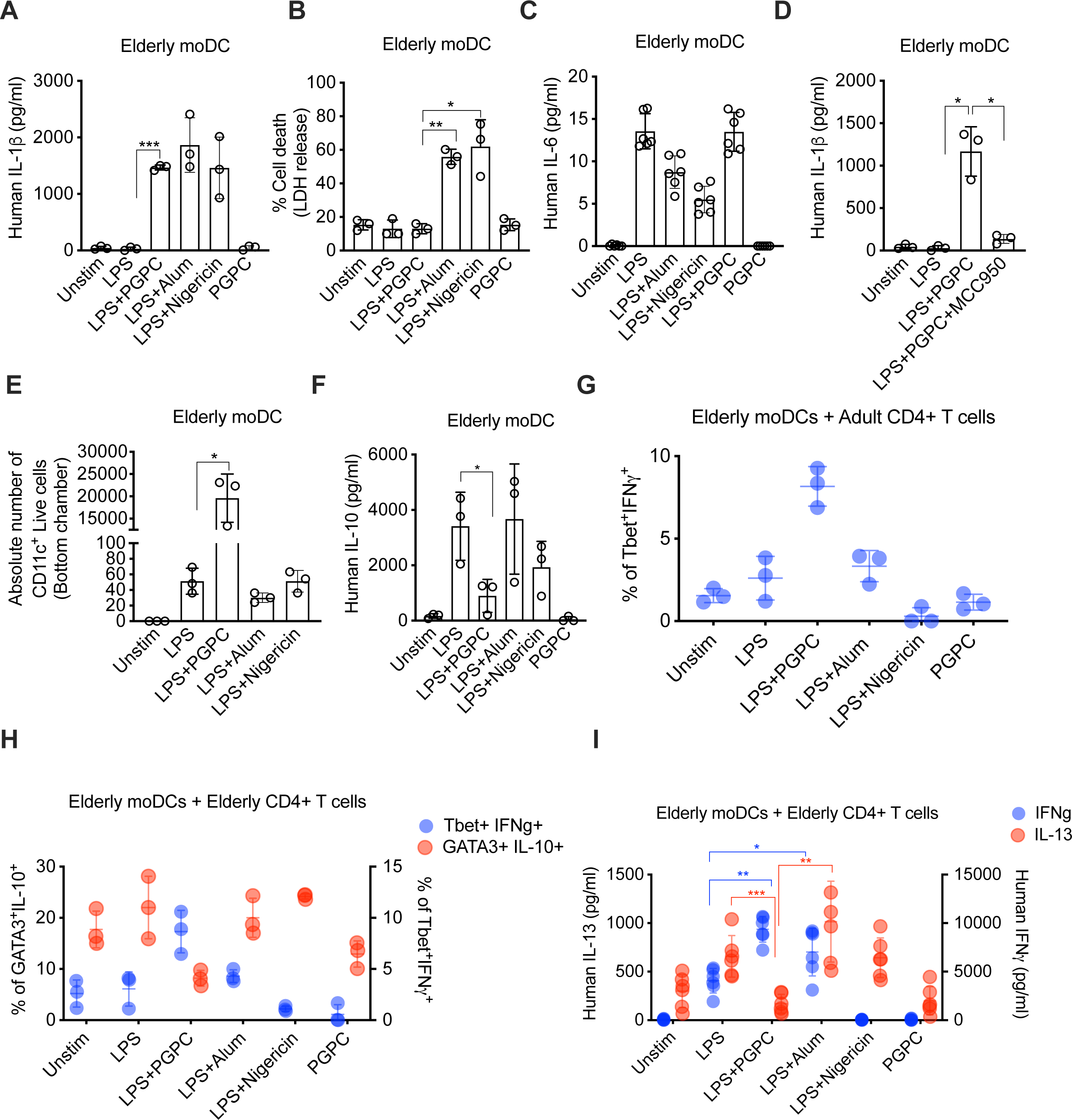
moDCs become hyperactive during aging and drive TH1-skewed immune responses. (A-D) Human moDC generated from elderly donors were either left untreated (‘‘Unstim’’) or treated with LPS alone, or alum alone, or PGPC alone for 24h, or DCs were primed for 3h with LPS, then treated with indicated stimuli for 21h. (A) IL-1β, (B) LDH release and (C) IL-6 were measured from 3 donors. (D) moDCs were either left untreated or treated with LPS, or moDCs were primed for 3h with LPS in the presence of the inhibitor MCC950, then treated with PGPC for 21h. IL-1β release from 3 donors was measured by ELISA. (E) moDCs were treated as indicated above. 24 hours post stimulation, moDCs were cultured in the upper chamber of a 24-well 5um transwell plate in the presence of CCL19. After 24h, the absolute number of cells that migrated to the bottom chamber was determined by flow cytometry. (F) moDCs were treated as above. IL-10 secretion was assessed by flow cytometry. (G) moDC from elderly donors were treated as above, then cultured for 5 days with allogeneic CD4^+^T cells from adult donors. The percentage of IFNγ^+^Tbet^+^ was calculated by flow cytometry. (H-I) moDC from elderly donors were treated as indicated, then cultured for 5 days with allogeneic CD4^+^T cells from elderly donors. The percentage of GATA3^+^IL-10^+^ and IFNγ^+^Tbet^+^ were measured by flow cytometry. (I) Human IFNγ and IL-13 secretion was measured by ELISA. Means and SDs of 3 donors is represented.

We next assessed if the hyperactivation of moDCs can improve moDC migratory activity, an attribute that is commonly defective in elderly DCs^2, 7^. moDCs from elderly individuals were stimulated with distinct stimuli and were subsequently cultured onto 5µm transwell plates in the presence of CCL19, a chemotactic ligand for CCR7. The absolute number of DCs that migrated to the bottom chamber was then calculated by flow cytometry. Naïve DC were not able to migrate in this assay. LPS induced the migration of few elderly moDCs, which is consistent with a dysfunction in moDC migration during aging^2, 7^. Notably, LPS+PGPC induced >250 fold more moDCs to migrate in this assay **(Figure 7E)**. moDC hyperactivators therefore correct age-associated defects in human moDC migration.

moDCs from the elderly secreted IL-10 when they were stimulated with LPS alone or LPS+alum or LPS+nigericin **(Figure 7F)**. IL-10 secretion was diminished when moDCs were stimulated with LPS+PGPC, as we have observed in murine DCs. These findings suggested that hyperactive moDCs may induce a skewed TH1 cells response, as IL-10 is a driver of TH2 responses. Using the MLR assay, we assessed the frequency of TH1 cells by intracellular staining using anti-Tbet and anti-IFNγ. We found that LPS+PGPC induced the highest frequency of Tbet^+^IFNγ^+^ among CD4^+^ T cells, as compared to moDCs activated with LPS or moDCs stimulated with LPS+alum or LPS+nigericin **(Figure 7G)**.

Finally, we tested the ability of hyperactive elderly moDCs to activate elderly T cells using the MLR assay. Unstimulated moDCs or moDCs stimulated with PGPC alone induced GATA3^+^IL-10^+^ TH2 cells but low IFNγ^+^Tbet^+^ cells **(Figure 7H-I)**. Conversely, pyroptotic stimuli enhanced GATA3^+^IL-10^+^ T cells but did not induce a higher frequency of Tbet^+^IFNγ^+^ TH1 cells **(Figure 7H-I)**. LPS+PGPC treated DCs did the opposite; they induced a higher frequency of IFNγ^+^Tbet^+^ but strongly diminished the frequency of GATA3^+^IL-10^+^ TH2 cells **(Figure 7H-I)**. Furthermore, cytokine release into the supernatants post co-culture confirmed that LPS+PGPC induced the highest levels of IFNγ secretion and conversely the lowest levels of IL-13 cytokine as compared to naïve, active or pyroptotic DCs **(Figure 7J)**. These data therefore establish that the hyperactivator LPS+PGPC enabled DCs to induce TH1 and suppress TH2 CD4^+^ T cell responses in elderly humans.

## Discussion

In this study, we explored age-associated changes in immune cell functions and corrected defects in DCs and T cells that normally undermine anti-tumor immunity. We found that while T cell targeted immunotherapies (*e.g.* anti-PD-1 or anti-CTLA4) eliminate tumors in young mice, these therapies are ineffective at eliminating the same tumors from elderly mice. This defect in the elderly immune system to response to checkpoint inhibitors tracks with our finding that naïve and total CD8^+^ T cells (which are often considered the target of PD-1 immunotherapies) are depleted with age. Similar findings have been made in humans, where the amounts of naïve CD8^+^ T cells diminish with age^56^. When combined with clinical data indicating that elderly humans are more susceptible to infection and cancer, this observed diminishment of CD8^+^ T cell functions is notable. We suggest that one of the reasons for the ineffectiveness of PD-1 based immunotherapies in the elderly is that these treatments are targeting the T cell subsets whose abundance is compromised in this age group. While these findings add to an increasingly long list of age-associated defects in immune cell functions, there are few examples of the ability to correct these defects. For example, age-associated defects in DC migration to dLNs could be restored to young DC levels by injecting animals with two times more elderly DCs^7^. However, increasing the amount of DCs injected into mice did not correct the age-associated impairment in anti-tumor immunity^7, 50^. This inability to correct functional defects in the elderly immune system frames the work performed herein.

We consider two central points of novelty associated with this study: 1) anti-tumor immunity in old mice is dependent on CD4^+^ T cells and not CD8^+^ T cells, and 2) stimuli that hyperactivate DCs can correct defects in immunity in old mice by stimulating robust CD4^+^ T cell responses. The decrease in CD8^+^ T cell importance with increased age is likely explained by their diminished abundance. The increased importance of CD4^+^ T cells observed with age, in contrast, remains unexplained, but similar findings have been made in long-term human cancer survivors who received CAR T cell treatments. CD4^+^ CAR T cells represent the dominant T cell population within cancer patients surviving 10+ years. In these studies, CD4^+^ CAR T cells exhibited cytotoxic characteristics long after remission^57–59^. In a glioblastoma orthotopic model, while CD8^+^ CAR T cells rapidly acquire an exhausted phenotype, the CD4^+^ CAR T cells persisted after tumor challenge and retained their effector potency^60^. The finding that different effector arms of the immune system are important at different stages of life undermines the value of single target (*i.e.* precision) immunotherapies. Conversely, this same finding highlights the value of targeting DCs directly, as these apex regulators of immunity are well-equipped to unleash multiple effector arms of immunity.

We found that only DC hyperactivators could induce antigen-specific T cell responses that persist over a year after immunization, and only DC hyperactivators induced T cell mediated anti-tumor immunity upon immunization of elderly mice. Our finding that CD4^+^ T cells from mice immunized over a year earlier could be transferred into naïve mice to confer anti-tumor immunity underscores the durability of the protective immunity generated by hyperactive DCs. The mechanism of this long-term immunity was linked to NLRP3 and IL-1β.

When combined with much prior work in this area, we can explain why hyperactive DCs are so effective at inducing durable T cell mediated immunity, and why other DC activation states are less effective. We propose that 5 signals are needed from DCs to maximally unleash their activities that drive age-agnostic T cell mediated protective immunity. These signals include: 1) MHC-based antigen presentation, 2) costimulatory molecule expression, 3) effector T cell skewing cytokines (*e.g.* IL-12p70), 4) memory T cell generating cytokines (*e.g.* IL-1β), and 5) high rates of DC migration to the dLN. The absence of any one of these signals would be expected to eliminate productive T cell stimulation by DCs, as we have found through the study of NLRP3-deficient cells (which cannot produce IL-1β) or CCR7-deficient cells (which cannot migrate). This idea also explains why adjuvants that induce DC activation (TLR ligands) or DC pyroptosis (alum) are of limited use in diverse clinical settings. Indeed, these stimuli are unable to induce all 5 of these activities, with TLR ligands unable to elicit IL-1β from DCs and unable to induce robust migration to the dLN. Similarly, it is well-recognized that alum induces TH2 immunity, rather than TH1 responses^27–29^. One reason for the lack of TH1-focused immunity of alum-treated cells is based on our findings that alum is a poor inducer of several of the 5 signals necessary for TH1 differentiation, such as CD40 expression, IL-12p70 secretion and cell migration. The lack of these activities likely renders other pyroptotic stimuli weak inducers of TH1 responses and anti-tumor protection. Based on the ability of DC hyperactivators to unleash all 5 of these activities from DCs, a wider range of optionality may be available by targeting hyperactive DCs.

Overall, our findings demonstrate that age-associated defects in immunity can not only be observed but can be functionally corrected. However, this correction has an interesting aspect. Rather than correcting the defects in naïve and total CD8^+^ T cell abundance in old mice, we found that protective immunity undergoes a shift from a dependence on CD8^+^ T cells to a dependence on CD4^+^ T cells. The immune system therefore has flexibility to use different effector arms, as conditions dictate, to serve the general purpose of host defense. These data provide a mandate to explore the mechanisms associated with age-induced changes in immunity, which may ultimately reveal a need to diversify the immune targets of therapeutic intervention.

## Materials and methods

### Mouse strains, and Tumor cell lines

C57BL/6J (Jax 000664), Balb/cJ (Jax 000651), *caspase-1/-113*^-/-^ mice (Jax 016621), *Nlrp3*^-/-^ KO (Jax 021302), *Ccr7*^-/-^ (Jax 006621), B6 Ly5.1 (Jax 002014), *Zbtb46*^DTR^ mice bearing the human diphtheria toxin receptor under zinc finger and BTB domain containing 46; also called zDC) gene (Jax 019506), and OT-II (Jax 004194) were purchased from Jackson Labs and used at 8 weeks old (young mice) or 68 weeks old (elderly mice) as indicated. For syngeneic tumor models in C57BL/6J, an OVA expressing cell line B16.F10OVA (a gift from Arlene Sharpe (Harvard)) was used. For a syngeneic colon cancer model in BALB/c mice, CT26 cell line was used (a gift from Jeff Karp (Harvard)).

### Reagents

*E. coli* LPS (Serotype O55:B5-TLR grade) was purchased from Enzo and used at 1μg/ml in cell culture or 10ug/mice for *in vivo* use. PGPC were purchased from Cayman Chemical. Reconstitution of commercially available PGPC was performed as described^52^. Briefly, ethanol solvent is evaporated using a gentle nitrogen gas stream. Pre-warmed serum-free media was then immediately added to the dried lipids to a final concentration of 1mg/ml. Reconstituted lipids were incubated at 37°C for 5-10 mins then sonicated for 20 s before adding to cells. PGPC were used at 100μg/ml or 50μg/ml for cell stimulation or 65μg/mice for *in vivo* use. EndoFit chicken egg ovalbumin protein with endotoxin levels <1 EU/mg was purchased from Invivogen for *in vivo* use at a concentration of 200μg/mice or *in vitro* use at a concentration ranging from 1000 to 10μg/ml. Incomplete Freund’s Adjuvant (F5506) was purchased from Sigma and used for *in vivo* immunizations at a working concentration of 1:2 (IFA:antigen emulsion). Alhydrogel referred to as alum was purchased from Accurate Chemical and used for in vitro stimulation at 100μg/ml and for *in vivo* immunization at a working concentration of 2mg/mouse. Nigericin was purchased from Invivogen and suspended in sterile ethanol to a stock concentration of 6.7 mM. Nigericin was used at a concentration of 10mM. MCC950 was purchased from invivogen and was used at 5uM.

### Murine Cell culture

FLT3-derived BMDCs were generated by differentiating bone marrow in IMDM (Gibco), 10% B16-FLT3L derived supernatant, 2μM 2-mercaptoethanol, 100 U/ml penicillin, 100μg/ml streptomycin (Sigma-Aldrich) and 10% FBS. 9 days after culture, BMDCs were washed with PBS and re-plated in IMDM with 10% FBS at a concentration of 1x10^6^ cells/ml in a final volume of 100μl. CD45R^neg^MHC-II^+^CD11c^+^DC purity was assessed by flow cytometry using BD Fortessa and was routinely above 80%. To induce active DCs, DCs were stimulated with LPS (1μg/ml) for 18 or 24 hours as indicated. To induce hyperactive or pyroptotic BMDCs, DCs were primed with LPS (1μg/ml) for 3 hours, then stimulated with PGPC (100μg/ml) or alum (100μg/ml) for 15h or 21h as indicated. T cells were cultured in RPMI-1640 (Gibco) supplemented with 10% FBS, 100 U/ml penicillin, 100μg/ml streptomycin (Sigma-Aldrich), and 50μM β-mercaptoethanol (Sigma-Aldrich). Tumor cell lines were all cultured in DMEM supplemented with 10% FBS. For OVA expressing cell lines, puromycin (2μg/ml) was added to the media.

### Human moDC generation and culture

Adult blood from healthy female and male volunteers aging between 25 and 45 years old, or elderly blood from female and male volunteers aging between 70 and 75years old were layered onto Ficoll-Hypaque and centrifuged. Mononuclear cells were recovered, and monocytes were then enriched using anti-Human CD14 beads (Miltenyi Biotec), according to the manufacturer’s instructions. Purity post enrichment was routinely measured by flow cytometry and measured above 85% of CD14 negative cells. Monocytes were cultured with 100 ng/mL (500 UI/mL) GM-CSF plus 50 ng/mL (250 UI/mL) IL-4 (Miltenyi Biotec) at 10^6^cells/mL in complete RPMI medium supplemented with 10% FCS (Gibco), 50 μM 2-mercaptoethanol, and 100 U/ml penicillin, 100μg/ml streptomycin (Sigma-Aldrich). 6 days post culture, monocyte derived dendritic cells (moDCs) were CD14^neg^CD11c^+^CD11b^+^CD1c^+^, and purity was greater than 92%. moDCs were cultured at 10^6^cells/mL in the presence of various stimuli to induce DC activation, hyperactivation or pyroptotosis. To induce DC activation, moDC were cultured with 1ug/mL of LPS alone. To induce DC hyperactivation or pyroptosis, moDCs were primed with LPS for 3 hours, then treated with PGPC (100ug/ml or 50ug/ml or 25ug/ml) or with alum (100ug/ml) or nigericin (10mM) for 7 hours or 21 hours as indicated. Supernatants were used to test cytokines by ELISA at indicated time points. To inhibit Gasdermin D oligomerization, necrosulfonamide (NSA) was used in culture 3 hours post priming. To inhibit NLRP3 oligomerization, MCC950 (5uM) was added to culture 3 hours post DC priming of LPS stimulation.

### Mixed leukocyte reaction

After 24 hours of moDCs stimulation, heterologous Human naive T cells from healthy volunteers were sorted as CD3^+^CD4^+^CD45RA^+^CD45RO^-^ and added to the culture at a ratio of 1:10 (DC:T cell) in the presence of IL-2 recombinant cytokine (50 ng/ml) (Peprotech). Five days post co-culture, supernatants were harvested for ELISA.

### Flow cytometry and cell sorting

After FcR blockade, 9 day old BMDCs were resuspended in MACS buffer (PBS with 1% FCS, 100 U/ml penicillin, 100μg/ml streptomycin and 2 mM EDTA) and stained for 20 minutes at 4 degrees with the following fluorescently conjugated antibodies (BioLegend): anti-CD11c (clone N418), anti-I-A/I-E (clone M5/114.15.2), anti-CD40 (clone 3/23), and anti-CCR7. Single cell suspension from the tumor or draining inguinal lymph nodes, or spleen were resuspended in MACS buffer and stained for 20 minutes at 4 degrees with the following fluorescently conjugated antibodies (BioLegend): anti-CD4 (clone RM4-5), anti-CD44 (clone IM7), anti-CD62L (clone MEL-14), anti-CD3 (clone 17A2), anti-CD45 (clone A20 or 30F11). LIVE/DEAD Fixable Violet or green Dead Cell Stain Kit (Molecular probes) was used to determine the viability of cells and cells were stained for 20 minutes in PBS at 4°C prior to antibodies staining. For antigen-specific T cell detection, T cells were stained with OVA-peptide tetramers at room temperature for 1h. APC conjugated I-A(b) AAHAEINEA (OVA 329-337) was used. I-A(b) associated with CLIP peptides were used as isotype controls. Tetramers were purchased for NIH tetramer core facility. To determine the absolute number of cells, countBright counting beads (Molecular probes) were used, following the manufacturer’s protocol. Appropriate isotype controls were used as a staining control. Data were acquired on a BD FACS ARIA or BD Fortessa (Becton-Dickenson). Data were analyzed using FlowJo software (Tree Star).

### LDH Assay and ELISA

Fresh supernatants were clarified by centrifugation then assayed for LDH release using the Pierce LDH cytotoxicity colorimetric assay kit (Life Technologies) following the manufacturer’s protocol. Measurements for absorbance readings were performed on a Tecan plate reader at wavelengths of 490 nm and 680 nm. To measure secreted cytokines, supernatants were collected, clarified by centrifugation, and stored at -20°C. ELISA for mouse IL-1β, TNFα, IL-10, IL-12p70, IFNγ, were performed using eBioscience Ready-SET-Go! (now ThermoFisher) ELISA kits according to the manufacturer’s protocol. ELISA for Human IL-1β and IL-10 were performed using ELISA MAX™ Deluxe Sets from Biolengend. Human IFNg and IL-13 cytokine secretion was measured by ELISA using eBioscience Ready-SET-Go! (now ThermoFisher) ELISA kits according to the manufacturer’s protocol.

### Migration Assay

Wild-type BMDCs generated using FLT3L were harvested on day 9 and suspended at a concentration of 5x10^6^ cells/ml in complete IMDM. DCs were cultured in polypropylene tubes with gentle rotation using MACSmix Tube Rotator (Milteny) in the incubator. DCs were either left untreated or treated with LPS alone for 24 hours, or BMDCs were primed with LPS for 3h then treated with PGPC or alum for 21h. Alternatively, BMDCs from *Nlrp3^-^*^/-^ or *Ccr7^-/-^* mice were primed with LPS for 3h then treated with PGPC for 21h. Cells were washed, stained for 30 minutes with CFSE, then 1x10^6^ DCs were injected subcutaneously on the right flank into ly5.1/CD45.1 mice in a total volume of 100ul. 12 hours post DC injection, single cell suspension from the skin draining lymph nodes was stained with live-dead violet kit in PBS, then stained in MACS buffer with anti-CD11c, anti-CD45.1 (clone A20) and anti-CD45.2 (clone 104) antibodies. Uninjected mice (no DCx) served as a control group. Hyperactive DCs that migrated to the dLN were sorted as CFSE^+^ CD45.2^+^ CD11c^+^ live cells, and resident DCs were sorted as CFSE^-^ CD45.1^+^ CD11c^+^ live cells, then cells were cultured in media alone for 24 hours onto 96-well round bottom plates. Supernatants were used for LDH and IL-1 cytokine release.

### Murine DC- T cell co-culture assay

Splenic CD4^+^ T cells were sorted from OT-II mice by magnetic cell enrichment using anti-CD4 beads (Milteny Biotech). T cells were then seeded in 96-well plates at a concentration of 10^5^ cells per well in the presence of 1x10^4^ FLT3L-DCs (10:1 ratio) that were generated from 8 weeks old mice and pretreated for 24 hours with either LPS alone or LPS+PGPC or LPS+alum and pulsed (or not) with OVA protein for 1 hour. For elderly T cells co-culture with DCs, splenic CD4^+^ T cells were sorted from 68 weeks old mice by magnetic cell enrichment using anti-CD4 beads (Milteny Biotech). T cells were then seeded in 96-well plates at a concentration of 10^5^ cells per well in the presence of 1x10^4^ FLT3L-DCs (10:1 ratio) that were generated from 8 weeks old mice or 68 weeks old mice and pretreated for 24 hours with either LPS alone or LPS+PGPC or LPS+alum and pulsed (or not) with OVA protein for 1 hour (500μg/ml) prior to co-culture. 4 days post culture, supernatants were collected and clarified by centrifugation for short-term storage at -20°C and cytokine measurement by ELISA.

### Generation of mixed bone marrow chimeric mice for immunizations

Four weeks old CD45.1^+^ female mice were exposed to whole body irradiation (2 doses of 500 rads per mouse, 2 hours apart). After at least 4 hours from the last irradiation, mice were reconstituted with 5x10^6^ bone marrow (BM) cells isolated from sex-matched mice and injected intravenously. Mice were kept in autoclaved cages and were treated with sulfatrim in the drinking water for 2 weeks after reconstitution. Mice were then placed in standard cages and allowed to reconstitute for 4 more weeks. To evaluate the percentage of chimerism, peripheral blood samples were collected at the end of reconstitution and stained for CD45.1 and CD45.2. For these experiments, all mice were housed at the BCH animal facility. To specifically deplete selected genes in cDCs, we generated mixed BM chimeras by reconstituting irradiated mice with 4x10^6^ BM cells isolated from *Zbtb46*^DTR^ mice mixed with 1x10^6^ BM cells isolated from wild-type, or Nlrp3^-/-^ mice. Reconstituted mice were then injected intraperitoneally with DTx (400 ng per mouse as first dose, then 200 ng per mouse) 3 times per week starting from 12 days prior to immunization for a total of 9 DTx dose.

### Generation of full bone marrow chimeric mice

To deplete conventional dendritic cells (cDCs) in aging mice, we generated full BM chimeras as mentioned above by reconstituting 60 weeks old irradiated mice with BM cells isolated from 60 weeks old *Zbtb46*^DTR^ mice. 6 weeks later, reconstituted mice were injected intraperitoneally with saline (control mice) or diphtheria toxin (DTx,400 ng per mouse) 3 times per week starting from 3 days post-tumor injection for a total of 6 DTx doses. DC depletion in the spleen and dLN was assessed 1-week post DT injection.

### *In vivo* immunization and T cell re-stimulation

Female C57BL/6J mice aged 8 weeks, were immunized subcutaneously (s.c.) on the right flank with either 200μg/mouse endotoxin-free OVA alone or with 10μg/mouse LPS emulsified in incomplete Freund’s adjuvant. Alternatively, wild type or *Nlrp3^-/-^*mice were immunized with 200μg/mouse endotoxin-free OVA, in combination with 10μg/mouse LPS plus 65μg/mouse PGPC as indicated, all emulsified in incomplete Freund’s adjuvant. In some experiments, mice were injected s.c. with OVA plus LPS emulsified in alum. When indicated, mice were s.c. injected with LPS and PGPC in the absence of OVA (NO OVA). 7 or 40 days after immunization, CD4^+^T cells were isolated from the skin draining lymph nodes of immunized mice by magnetic cell sorting with anti-CD4 microbeads (Miltenyi). Enriched cells were then sorted as live CD45^+^CD3^+^CD4^+^. Purity post-sorting was >98%. Sorted cells were then seeded in 96-well plates in the presence of OVA-preloaded BMDC (at a ratio of 1 DC:10 T cells). In some experiments, 8 week old mice were immunized s.c with OVA (200μg/mouse) alone emulsified in IFA. 7 days later, mice received a s.c. boost with OVA protein (200μg/mouse) alone emulsified in IFA. One week after OVA boost, memory CD4^+^T cells from the skin dLN were enriched by positive selection using naïve CD4+ T cell kit containing anti-CD44 microbeads (Miltenyi). CD44^+^CD4^+^ T cells were sorted and restimulated in vitro with OVA-preloaded BMDC (at a ratio of 1 DC:10 T cells). In other experiments, 8 weeks old mice were with OVA (200μg/mouse) alone emulsified in IFA, mice received a s.c. boost with OVA protein (200μg/mouse) alone emulsified in IFA. 60 weeks later, mice received a s.c. boost with OVA alone (200μg/mouse) emulsified in IFA. One week after OVA re-exposure, CD4^+^T cells were sorted from the dLN and restimulated in vitro with OVA-preloaded BMDC (at a ratio of 1 DC:10 T cells). When indicated, 68 weeks old mice were with immunized with OVA (200μg/mouse) alone emulsified in IFA. One week later, mice received a s.c. boost with OVA protein (200μg/mouse) alone emulsified in IFA. One week after OVA re-exposure, CD4^+^T cells were sorted from the dLN and restimulated in vitro with OVA-preloaded BMDC (at a ratio of 1 DC:10 T cells). Secretion of IFNγ, IL-4 or IL-10 was measured by ELISA 5 days later. In some experiments, the percentage of antigen-specific T cells expressing IFNγ was measured by using AAHAEINEA tetramer staining as AAHAEINEA^+^CD4^+^CD3^+^ T cells followed by an IFNγ intracellular staining 5 days post-co-culture.

### Intracellular staining

For intracellular cytokine staining, T cells were stimulated with 50 ng/ml phorbol 12-myristate 13-acetate (PMA) and 500 ng/ml ionomycin (Sigma-Aldrich) in the presence of GolgiStop (BD) and brefeldin A for 4 h. Cells were then washed twice with PBS, and stained with LIVE/DEAD Fixable violet Stain Kit (Molecular probes) in PBS for 20 min at 4°C. T cells were then washed with MACS buffer, and stained for appropriate surface markers for 20 min at 4°C. After two washes, cells were fixed and permeabilizated using BD Cytofix/Cytoperm kit for 20 min at 4°C, then washed with 1X perm wash buffer (BD) per manufacturer’s protocol. Intracellular cytokine staining was performed in 1X perm buffer for 20-30 min at 4°C using the following conjugated antibodies all purchased from BioLegend: mouse anti-IFN-γ (clone XMG1.2), mouse anti-TNFα (clone MP6-XT22), anti-mouse IL-10 (clone JES5-16E3), anti-mouse IL4 (clone 11B11), GATA3 (clone 16E10A23), T-bet (clone 4B10). For Tregs staining, True-Nuclear™ One Step Staining Mouse Treg Flow™ Kit (FOXP3 (clone MF-14) with CD25 and CD4) was used with FOXP3 Fix/Perm Buffer from BioLegend per manufacturer’s protocol. Data were acquired on a BD FACS ARIA or BD Fortessa. Data were analyzed using FlowJo software.

### Whole tumor lysates preparation

To prepare whole tumor cell lysates (WTL) syngeneic from tumors explants, tumors from unimmunized mice bearing a tumor 10-12 mm of size were mechanically disaggregated using gentleMACS dissociator (Milteny Biotec), and digested using the tumor Dissociation Kit (Milteny Biotec) following the manufacturer’s protocol. Tumors were incubated for 45 minutes at 37 degrees in a tube rotator for complete digestion. After digestion, tumors were washed with PBS and passed through 70-μm and 30-μm filters. Single cell suspension were depleted of CD45^+^ cells using anti-CD45 TILs microbeads (Milteny Biotec). Tumor cells were then counted and resuspended at 5x10^6^ cells/ml then lysed by 3-4 cycles of freeze-thawing. Lysates were sheered using a syringe and 18G then 21G needle until the lysate is clear of cell clumps. Lysates were then centrifuged at 12,000 rpm for 15 minutes to remove all cellular debris, passed through 70-μm and 30- μm filters then stored in aliquots at -20°C until use. WTL were used for immunotherapy at a concentration equivalent to 2.5x10^5^ tumor cells per mice.

### Prophylactic immunization and tumor challenges

For immunizations prior to tumor inoculation, 8 weeks old Female C57BL/6J mice were injected subcutaneously (s.c.) into the right flank with OVA (200μg/mouse) alone (PBS), or in combination with 10μg/mice of LPS, or with 10μg/mice of LPS plus 65μg/mice of PGPC, all emulsified in incomplete Freud’s adjuvant (IFA). Some mice were injected s.c. with OVA plus LPS emulsified in alum. Alternatively, 8 weeks old *Nlrp3^-/-^* mice were s.c. immunized with 10μg/mice of LPS plus 65μg/mice of PGPC emulsified in IFA. 60 weeks post immunization, all mice received a s.c. boost with OVA protein (200μg/mouse) alone emulsified in IFA. One week later, all mice were challenged s.c. on the left flank with 3x10^5^ of viable B16OVA cells. The size of the tumors was assessed in a blinded, coded fashion every two days and recorded as tumor area (length × width) using a caliper. Mice were sacrificed when tumors reached 2 cm^3^ or upon ulceration.

### Immunotherapeutic immunization and tumor challenges

For immunizations in the context of an immunotherapeutic approach, 8 weeks or 68 weeks old C57BL/6J were injected on the left flank with 3x10^5^ of viable B16OVA cells. Alternatively, BALB/c mice were injected on the left flank with 3x10^5^ viable CT26 cells. When tumors reached 2-3mm of size, mice were either left untreated (unimmunized) or mice were immunized with WTL alone or in combination with 10μg/mice of LPS and 65μg/mice of PGPC emulsified in incomplete Freud’s adjuvant (IFA). In some experiment, tumor bearing mice were immunized with WTL and 10μg/mice of LPS plus 2mg/mouse of alum. Immunizations were followed by two boost injections every 7-10 days. Immunized mice were divided blindly into several groups. Some mice were injected intraperitoneally (i.p.) with 100μg of Ultra-LEAF anti-CD4 (clone GK1.5), or CD8a (clone 53-6.7) one day prior the day of immunization, followed by 5 consecutive injections every 2 days. Other mice were given 100μg of LEAF anti-mouse/rat IL-1β antibody by i.v. injection every day for four consecutive days; starting two days before receiving the immunization, then on day 1, day 2, post-immunization to ensure chronic depletion of circulating IL-1β. The injection of anti-IL-1β antibody was repeated for every boost injection. Control mice received isotype-matched rat IgG. In some experiments, when tumors reached 2-3mm of size, mice were either left untreated (unimmunized) or injected i.p. with 100μg of anti-PD-1 or 100μg of anti-CTLA4 or a combination of 100μg anti-CTL4 and 100μg of anti-PD1 every two days for 5 consecutive injections. All antibodies were purchased from BioLegend. The size of the tumors was assessed in a blinded, coded fashion every two days and recorded as tumor area (length × width) using a caliper. Mice were sacrificed when tumors reached 2 cm^3^ or upon ulceration.

### Tumor infiltrating Tregs and TILs activation

To assess the frequency of T regulatory cells (Tregs) among tumor-infiltrating lymphocytes (TIL), tumors from unimmunized or mice immunized with WTL+LPS+PGPC were harvested then dissociated using the tumor Dissociation Kit and the gentleMACS dissociator (Miltenyi Biotec), following the manufacturer’s protocol. After digestion, tumors were washed with PBS and passed through 70- μm and 30-μm filters. CD45^+^ cells were positively selected using anti-CD45 TILs microbeads (Miltenyi Biotec). The frequency of Tregs infiltrating in the tumor microenvironment was defined as FOXP3^+^CD25^+^CD4^+^ among CD45^+^ live cells and assessed by flow cytometry. Tumor infiltrating CD45^+^ cells were cultured for 24 hours with dynabeads mouse T-Activator CD3/CD28 (Gibco) for T cell activation, followed by 5 hours stimulation with PMA and ionomycin. IFN-γ and IL-10 cytokine expression by CD4^+^ TILs was assessed by Intracellular staining.

### Adoptive cell transfer

8 weeks old mice were immunized with B16OVA WTL and LPS plus PGPC emulsified in in IFA. 60 weeks later, mice received a s.c. boost with OVA emulsified in IFA. One week after OVA re-exposure, CD4^+^, CD8^+^ T cells, CD19^+^ B cells, and CD14^+^ monocytes were enriched from the spleen using anti CD4, anti-CD8, anti-CD19 or anti-CD14 microbeads (Miltenyi Biotec). Enriched cells were then sorted as live CD45^+^ CD3^+^ CD4^+^ or CD45^+^ CD3^+^ CD8^+^ or CD45^+^ CD3^neg^ CD19^+^ or CD45^+^ CD3^neg^ CD14^+^ using FACS ARIA.

Purity post-sorting was >95%. Sorted CD4^+^, CD8^+^ T cells were then stimulated for 24 h in 24-well plates (∼2×10^6^ cells/well) coated with anti-CD3 (4 μg/ml) and anti-CD28 (4 μg/ml) in the presence of IL-2 (Peprotech, 50ng/ml). CD19^+^ B cells were stimulated for 24 hours using Purified F(ab’)2 Goat anti-mouse IgM Antibody (BioLegend). CD14^+^ monocytes were stimulated for 24 hours with 1ug/ml of LPS. 5×10^5^ of activated CD4^+^cells, or CD19^+^, or CD14^+^, and 2×10^5^ of activated CD8+ T cells were then transferred by i.v. injection on the right flank into naïve recipient 8 weeks old mice. 7 days post transfer, all mice were s.c. challenged with 3×10^5^ of viable B16OVA cells. The size of the tumors was assessed in a blinded, coded fashion every two days and recorded as tumor area (length × width) using a caliper. Mice were sacrificed when tumors reached 2 cm^3^ or upon ulceration.

### Statistical analysis

Statistical significance for experiments with more than two groups was tested with two-way ANOVA with Tukey multiple comparison test correction. Kaplan Meier (Log-rank Mantel cox) test was used to compare the survival of tumor bearing mice. Adjusted p-values, calculated with Prism (Graphpad), are coded by asterisks: <0.05 (*); < 0.01(**); <0.0005 (***); ≤0.0001 (****).

## Supplementary Figures

**Figure S1.**
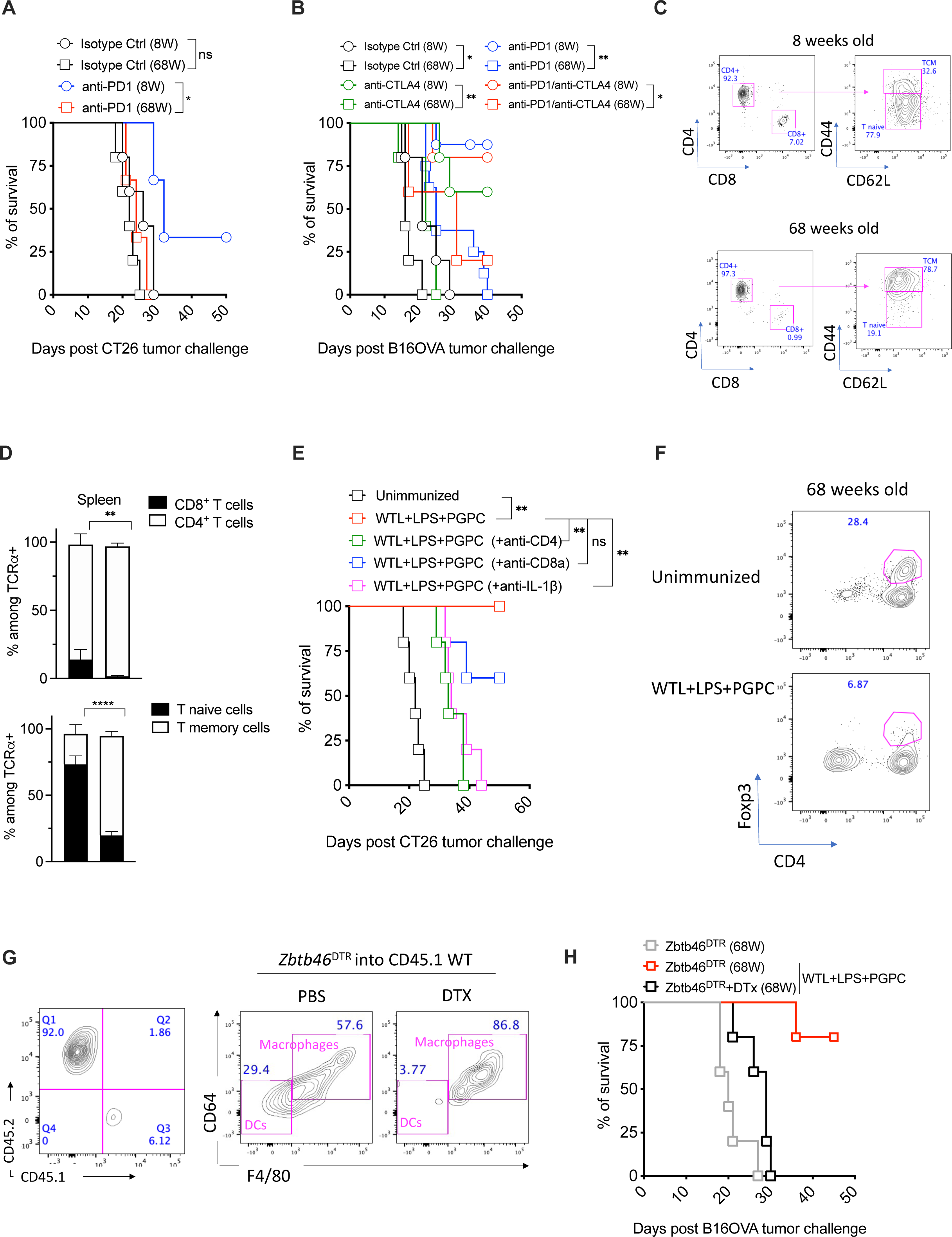
CD4^+^ T cells mediate anti-tumor immunity in young and old mice. 8 weeks or 68 weeks old Balbc/J mice were injected subcutaneously (s.c.) on the right flank with CT26. When tumors reached 2 mm of size, mice were either left untreated (unimmunized) or treated i.p. with anti-PD1 antibodies. The percentage of mice survival was monitored every 2 days (n=5 mice per group). (B) C57BL/6J mice were injected with B16OVA cells. 7 days post injection, B16OVA tumor-bearing mice were either treated with anti-PD1 antibodies alone or anti-CTLA4 alone or a combination of anti-PD1 and anti-CTLA4 antibodies for 5 consecutive injections. Tumor size in mm^2^ was measured every two-three days (n=5 mice per group). (C) Representative plots and (D) histograms indicating the percentage of CD4^+^ and CD8^+^ T cells among splenic TCRa^+^ CD45^+^ cells and the percentages of naïve and memory T cells as measured by flow cytometry in 8 weeks or 68 weeks old C57BL/6J mice. (E) 68 weeks old Balbc/J mice were injected s.c. on the right flank with CT26 cells. When tumors reached 2-3mm in size, mice were either left unimmunized or were immunized with LPS plus PGPC with or without neutralizing anti-CD4, or anti-IL-1β or anti-CD8a antibodies. Percentage of mice survival was monitored every 2 days (n=5 mice per group). (G-H) 68 weeks old mice on a CD45.1 background were irradiated and reconstituted with *Zbtb46*^DTR^ bone marrow. 6 weeks post-reconstitution, the efficiency of reconstitution was measured by flow cytometry (left panel). *Zbtb46*^DTR^ chimeric mice were injected with diphteria toxin (DTx) 3 times a week for a total of 9 DTx injections. The efficiency of conventional DC depletion was measured by flow cytometry (right panel). 6 weeks post-reconstitution, *Zbtb46*^DTR^ chimeric mice were s.c. injected on the right flank with B16OVA cells. (H) When tumors reached 2-3 mm of size mice were injected or not with DTx 3 times a week for a total of 9 DTx injections, and either left unimmunized or were immunized with WTL and LPS plus PGPC. The percentage of mice survival was monitored every 2 days (n=5 mice per group).

**Figure S2.**
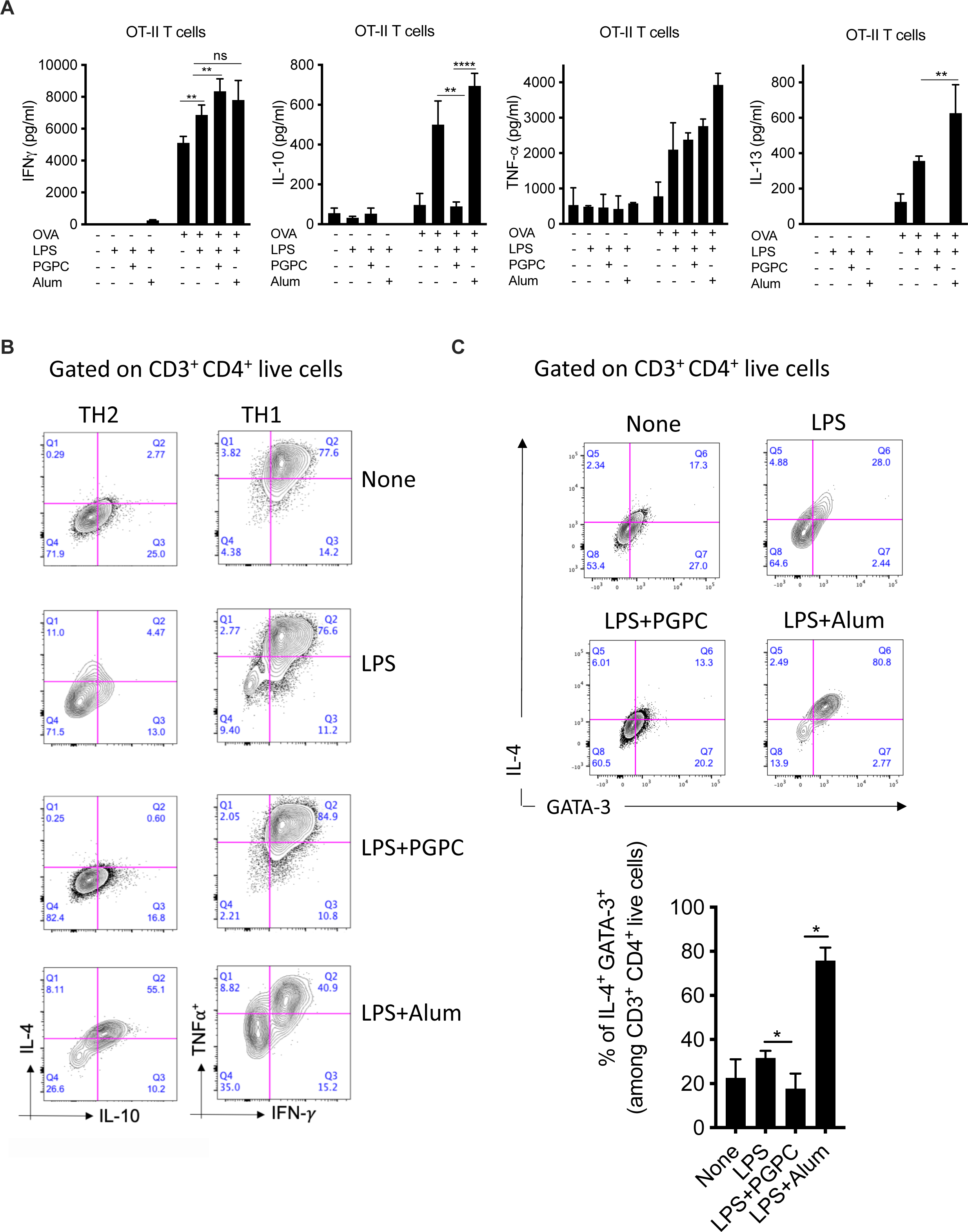
Hyperactive DCs drive TH1-skewed immune responses, with no evidence of TH2 immunity. (A-C) BMDCs pretreated with indicated stimuli were loaded (or not) with OVA protein for 1h, then incubated for 4 days with OT-II naïve CD4+ T cells. (A) Supernatants were collected and IFNγ, IL-10, TNFa and IL-13 cytokine release was measured by ELISA. (B-C) T cells were stimulated for 5h with PMA plus ionomycin in the presence of brefeldin-A and monensin. (B) Representative plots of the frequency of TH1 cells as TNFa^+^ IFNg^+^, and TH2 cells as IL-4^+^IL-10^+^ and (C) The frequency of GATA3^+^ IL-4^+^ TH2 cells were measured following intracellular staining by flow cytometry. Histograms represent the frequency of IL-10^+^GATA3^+^. Means and SDs of five mice are shown.

**Figure S3.**
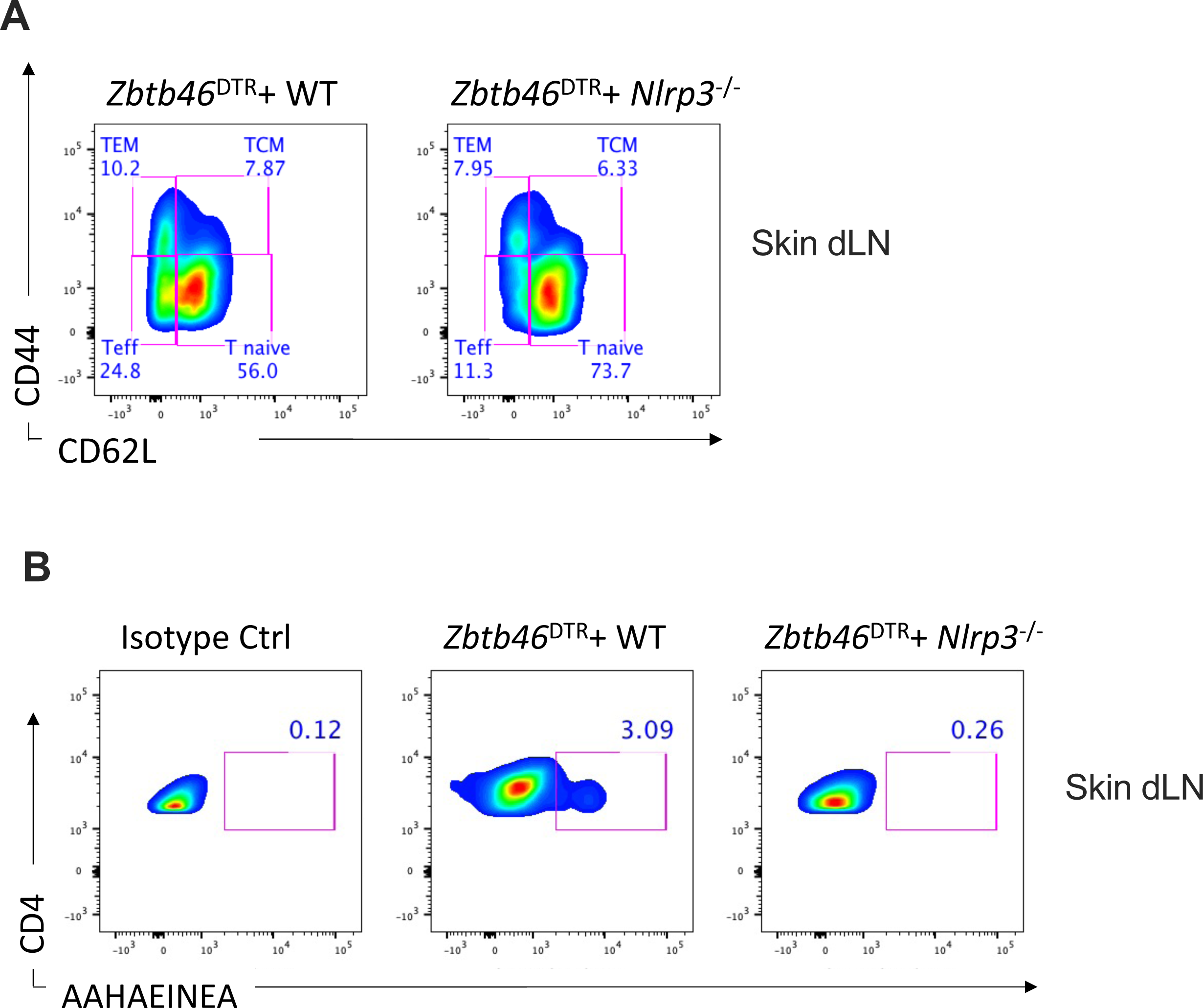
DC hyperactivators enhance CD4^+^ reactivation of already committed TH1 cells and accelerate memory CD4^+^T cell generation in an inflammasome dependent manner. (A-B) CD45.1 mice were irradiated then reconstituted with mixed Bone marrow of the genotypes indicated. Six weeks post-reconstitution, chimera mice were injected with diphteria toxin (DTx) 3 times a week for a total of 9 DTx injections. Chimeric mice were then immunized s.c. on the right flank with OVA with LPS plus PGPC. Seven days post-immunization with OVA with LPS and PGPC. (A) Representative plot of the percentage of Teff, TEM, TCM, and T naive cells in the skin dLN as measured by flow cytometry. (B) Representative plot of the percentage of AAHAEINEA^+^ among CD4^+^ live T cells from the dLN as measured using OVA peptide tetramer staining. Means and SDs of five mice are shown.

**Figure S4.**
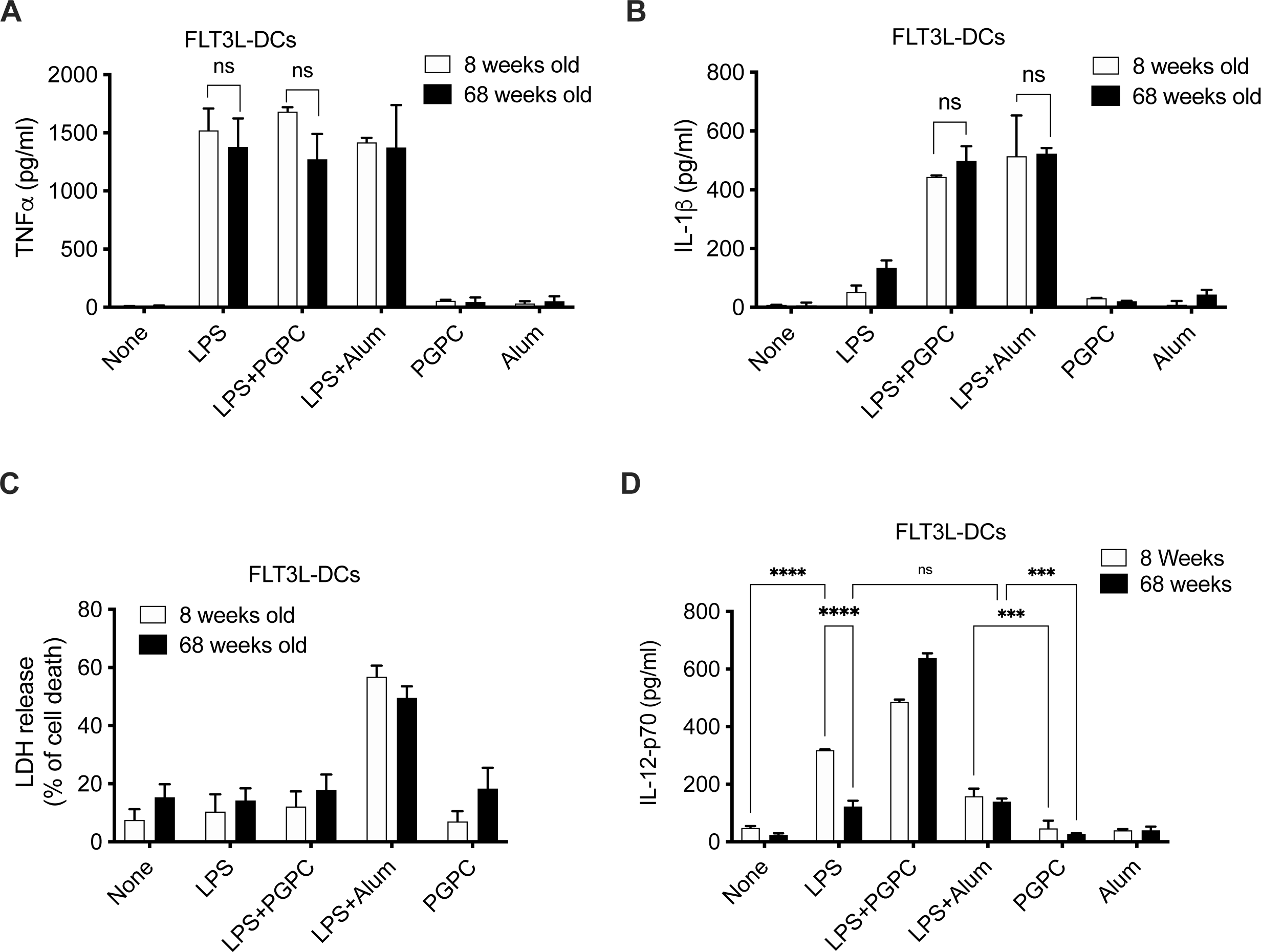
DCs from elderly mice achieve a state of hyperactivation. (A-D) 8 weeks or 68 weeks old FLT3L-derived BMDCs were either left untreated (‘‘None’’) or treated with LPS alone, or alum alone, or PGPC alone for 24h, or BMDCs were primed for 3h with LPS, then treated with indicated stimuli for 21h. (A) TNFa and (B) IL-1β release was monitored by ELISA. (C) LDH release by FLT3L-derived BMDCs was measured. (D) BMDCs were treated as described in A and cultured for 24 hours onto an anti-CD40 coated plate. IL-12p70 release was monitored by ELISA. Means and SDs from three replicates are shown, and data are representative of at least three independent experiments.

**Figure S5:**
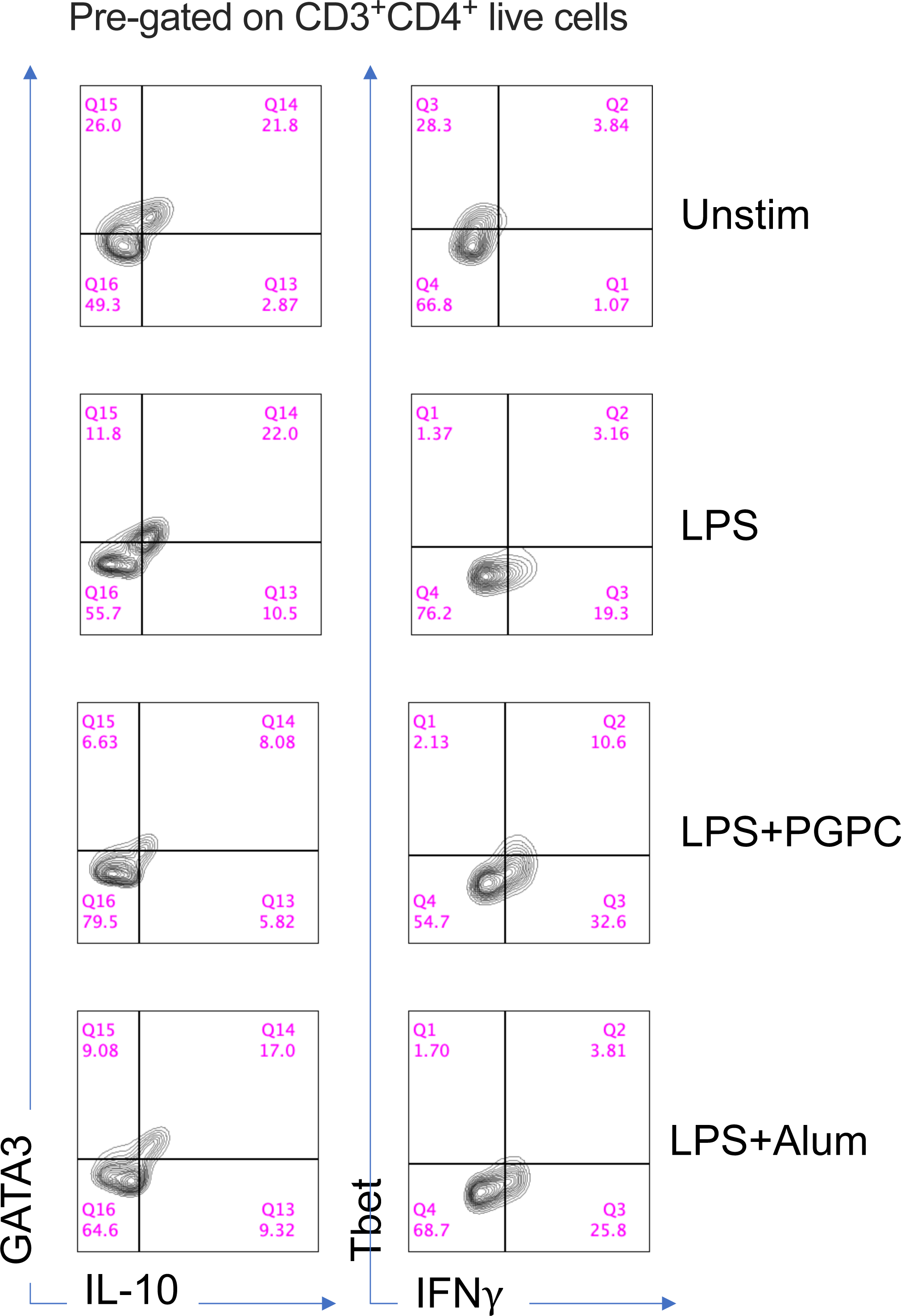
moDCs drive TH1-skewed immune responses. Human moDC generated from elderly donors were either left untreated (‘‘Unstim’’) or treated with LPS alone, or alum alone, or PGPC alone for 24h, or DCs were primed for 3h with LPS, then treated with indicated stimuli for 21h. Treated DCs were then cultured for 5 days with allogeneic CD4^+^T cells from elderly donors. Representative plots indicate the percentage of GATA3^+^IL-10^+^ and IFNγ^+^Tbet^+^ among CD4+ T cells were measured by flow cytometry.

## References

1. Akbar, A. N. & Gilroy, D. W. Aging immunity may exacerbate COVID-19. Science *(1979)* 369, 256–257 (2020).

2. Agrawal, A. & Gupta, S. Impact of aging on dendritic cell functions in humans. Ageing Research Reviews vol. 10 336–345 Preprint at https://doi.org/10.1016/j.arr.2010.06.004 (2011).

3. Bailey, K. L. et al. Aging leads to dysfunctional innate immune responses to TLR2 and TLR4 agonists. Aging Clin Exp Res 31, 1185–1193 (2019).

4. Montgomery, R. R. & Shaw, A. C. Paradoxical changes in innate immunity in aging: recent progress and new directions. J Leukoc Biol 98, 937–943 (2015).

5. Agrawal, A. et al. Altered Innate Immune Functioning of Dendritic Cells in Elderly Humans: A Role of Phosphoinositide 3-Kinase-Signaling Pathway. The Journal of Immunology 178, 6912– 6922 (2007).

6. Pereira, L. F., Duarte de Souza, A. P., Borges, T. J. & Bonorino, C. Impaired in vivo CD4+ T cell expansion and differentiation in aged mice is not solely due to T cell defects: decreased stimulation by aged dendritic cells. Mech Ageing Dev 132, 187–194 (2011).

7. Grolleau-Julius, A., Harning, E. K., Abernathy, L. M. & Yung, R. L. Impaired Dendritic Cell Function in Aging Leads to Defective Antitumor Immunity. Cancer Res 68, 6341 (2008).

8. Ciabattini, A. et al. Vaccination in the elderly: The challenge of immune changes with aging. Seminars in Immunology vol. 40 83–94 Preprint at https://doi.org/10.1016/j.smim.2018.10.010 (2018).

9. Gravekamp, C. & Chandra, D. Aging and cancer vaccines. Crit Rev Oncog 18, 585–95 (2013).

10. Pinti, M. et al. Aging of the immune system: Focus on inflammation and vaccination. European Journal of Immunology vol. 46 2286–2301 Preprint at https://doi.org/10.1002/eji.201546178 (2016).

11. Park, J. A. & Cheung, N. K. V. Limitations and opportunities for immune checkpoint inhibitors in pediatric malignancies. Cancer Treat Rev 58, 22–33 (2017).

12. Naltet, C. & Besse, B. Immune checkpoint inhibitors in elderly patients treated for a lung cancer: a narrative review. Transl Lung Cancer Res 10, 3014–3028 (2021).

13. Abdel-Rahman, O. et al. Treatment-related Death in Cancer Patients Treated with Immune Checkpoint Inhibitors: A Systematic Review and Meta-analysis. Clin Oncol (R Coll Radiol) 29, 218–230 (2017).

14. Fitzgerald, K. A. & Kagan, J. C. Toll-like Receptors and the Control of Immunity. Cell vol. 180 1044–1066 Preprint at https://doi.org/10.1016/j.cell.2020.02.041 (2020).

15. Alloatti, A., Kotsias, F., Magalhaes, J. G. & Amigorena, S. Dendritic cell maturation and cross-presentation: timing matters! Immunological Reviews vol. 272 97–108 Preprint at https://doi.org/10.1111/imr.12432 (2016).

16. Iwasaki, A. & Medzhitov, R. Control of adaptive immunity by the innate immune system. Nat Immunol 16, 343–53 (2015).

17. Alvarez, D., Vollmann, E. H. & von Andrian, U. H. Mechanisms and consequences of dendritic cell migration. Immunity 29, 325–42 (2008).

18. van den Eeckhout, B., Tavernier, J. & Gerlo, S. Interleukin-1 as Innate Mediator of T Cell Immunity. Frontiers in immunology vol. 11 621931 Preprint at https://doi.org/10.3389/fimmu.2020.621931 (2020).

19. Ben-Sasson, S. Z., Wang, K., Cohen, J. & Paul, W. E. IL-1 Strikingly Enhances Antigen-Driven CD4 and CD8 T-Cell Responses. Cold Spring Harb Symp Quant Biol 78, 117–124 (2013).

20. Ben-Sasson, S. Z. et al. IL-1 acts directly on CD4 T cells to enhance their antigen-driven expansion and differentiation. Proc Natl Acad Sci U S A 106, 7119–24 (2009).

21. Ben-Sasson, S. Z. et al. IL-1 enhances expansion, effector function, tissue localization, and memory response of antigen-specific CD8 T cells. J Exp Med 210, 491–502 (2013).

22. Lee, P.-H. et al. Host conditioning with IL-1β improves the antitumor function of adoptively transferred T cells. J Exp Med jem.20181218 (2019) doi:10.1084/jem.20181218.

23. Sarkar, S. et al. Programming of CD8 T Cell Quantity and Polyfunctionality by Direct IL-1 Signals. J Immunol 201, 3641–3650 (2018).

24. Jain, A., Song, R., Wakeland, E. K. & Pasare, C. T cell-intrinsic IL-1R signaling licenses effector cytokine production by memory CD4 T cells. Nat Commun 9, 1–13 (2018).

25. Evavold, C. L. & Kagan, J. C. Inflammasomes: Threat-Assessment Organelles of the Innate Immune System. Immunity 51, 609–624 (2019).

26. Martinon, F., Burns, K. & Tschopp, J. The Inflammasome: A molecular platform triggering activation of inflammatory caspases and processing of proIL-β. Mol Cell 10, 417–426 (2002).

27. Oleszycka, E. et al. The vaccine adjuvant alum promotes IL-10 production that suppresses Th1 responses. Eur J Immunol 48, 705–715 (2018).

28. Brewer, J. M. et al. Aluminium hydroxide adjuvant initiates strong antigen-specific Th2 responses in the absence of IL-4- or IL-13-mediated signaling. J Immunol 163, 6448–54 (1999).

29. Mori, A. et al. The vaccine adjuvant alum inhibits IL-12 by promoting PI3 kinase signaling while chitosan does not inhibit IL-12 and enhances Th1 and Th17 responses. Eur J Immunol 42, 2709–2719 (2012).

30. Zhivaki, D. et al. Inflammasomes within Hyperactive Murine Dendritic Cells Stimulate Long-Lived T Cell-Mediated Anti-tumor Immunity. CellReports 33, 108381 (2020).

31. Nikolich-Žugich, J. The twilight of immunity: emerging concepts in aging of the immune system. Nat Immunol 19, 10–19 (2018).

32. Wang, Y. et al. Combination of Fruquintinib and Anti–PD-1 for the Treatment of Colorectal Cancer. The Journal of Immunology 205, 2905–2915 (2020).

33. Garris, C. S. et al. Successful Anti-PD-1 Cancer Immunotherapy Requires T Cell-Dendritic Cell Crosstalk Involving the Cytokines IFN-γ and IL-12. Immunity 49, 1148–1161.e7 (2018).

34. Marrack, P., McKee, A. S. & Munks, M. W. Towards an understanding of the adjuvant action of aluminium. Nat Rev Immunol 9, 287–293 (2009).

35. Garçon, N. & di Pasquale, A. From discovery to licensure, the Adjuvant System story. Hum Vaccin Immunother 13, 19–33 (2017).

36. Robert, C. A decade of immune-checkpoint inhibitors in cancer therapy. Nat. Commun. 11, 3801 (2020).

37. Sharpe, A. H. & Pauken, K. E. The diverse functions of the PD1 inhibitory pathway. Nat. Rev. Immunol. 18, 153–167 (2018).

38. Martins, C. et al. Distinct antibody clones detect PD-1 checkpoint expression and block PD-L1 interactions on live murine melanoma cells. Scientific Reports 2022 12:*1* 12, 1–14 (2022).

39. Goronzy, J. J. & Weyand, C. M. Successful and Maladaptive T Cell Aging. Immunity vol. 46 364–378 Preprint at https://doi.org/10.1016/j.immuni.2017.03.010 (2017).

40. Zanoni, I. et al. An endogenous caspase-11 ligand elicits interleukin-1 release from living dendritic cells. Science *(1979)* 352, 1232–1236 (2016).

41. Newman, M. J. et al. Saponin adjuvant induction of ovalbumin-specific CD8+ cytotoxic T lymphocyte responses. J Immunol 148, 2357–62 (1992).

42. Kool, M. et al. Alum adjuvant boosts adaptive immunity by inducing uric acid and activating inflammatory dendritic cells. J Exp Med 205, 869–882 (2008).

43. Marichal, T. et al. DNA released from dying host cells mediates aluminum adjuvant activity. Nat Med 17, 996–1002 (2011).

44. Barnden, M. J., Allison, J., Heath, W. R. & Carbone, F. R. Defective TCR expression in transgenic mice constructed using cDNA-based α- and β-chain genes under the control of heterologous regulatory elements. Immunol Cell Biol 76, 34–40 (1998).

45. Sallusto, F. & Lanzavecchia, A. Heterogeneity of CD4+ memory T cells: functional modules for tailored immunity. Eur J Immunol 39, 2076–82 (2009).

46. Lanzavecchia, A. & Sallusto, F. Understanding the generation and function of memory T cell subsets. Curr Opin Immunol 17, 326–332 (2005).

47. Ben-Sasson, S. Z. et al. IL-1 enhances expansion, effector function, tissue localization, and memory response of antigen-specific CD8 T cells. J Exp Med 210, 491–502 (2013).

48. Zanoni, I. et al. IL-15 cis Presentation Is Required for Optimal NK Cell Activation in Lipopolysaccharide-Mediated Inflammatory Conditions. Cell Rep 4, 1235–1249 (2013).

49. Moro-García, M. A., Alonso-Arias, R. & Lopez-Larrea, C. When Aging Reaches CD4+ T-Cells: Phenotypic and Functional Changes. Front Immunol 0, 107 (2013).

50. Grolleau-Julius, A., Abernathy, L., Harning, E. & Yung, R. L. Mechanisms of murine dendritic cell antitumor dysfunction in aging. Cancer Immunol Immunother 58, 1935–9 (2009).

51. Grolleau-Julius, A., Harning, E. K., Abernathy, L. M. & Yung, R. L. Impaired Dendritic Cell Function in Aging Leads to Defective Antitumor Immunity. Cancer Res 68, 6341 (2008).

52. Evavold, C. L. et al. The Pore-Forming Protein Gasdermin D Regulates Interleukin-1 Secretion from Living Macrophages. Immunity 48, 35–44.e6 (2018).

53. Rathkey, J. K. et al. Chemical disruption of the pyroptotic pore-forming protein gasdermin D inhibits inflammatory cell death and sepsis. Sci Immunol 3, 2738 (2018).

54. Coll, R. C. et al. MCC950 directly targets the NLRP3 ATP-hydrolysis motif for inflammasome inhibition. Nature Chemical Biology 2019 15:*6* 15, 556–559 (2019).

55. Tapia-Abellán, A. et al. MCC950 closes the active conformation of NLRP3 to an inactive state. Nature Chemical Biology 2019 15:*6* 15, 560–564 (2019).

56. Nikolich-Žugich, J., Li, G., Uhrlaub, J. L., Renkema, K. R. & Smithey, M. J. Age-Related Changes in CD8 T Cell Homeostasis and Immunity to Infection. Semin Immunol 24, 356 (2012).

57. Melenhorst, J. J. et al. Decade-long leukaemia remissions with persistence of CD4+ CAR T cells. Nature 602, 503–509 (2022).

58. Flemming, A. Cytotoxic CD4+ CAR T cells implicated in long-term leukaemia remission. Nature Reviews Immunology 2022 22:*3* 22, 146–146 (2022).

59. Chen, A. X. Y., Derrick, E. B., Beavis, P. A. & House, I. G. CD4+ chimeric antigen receptor T cells in for the long journey. Immunol Cell Biol 100, 304–307 (2022).

60. Wang, D. et al. Glioblastoma-targeted CD4+ CAR T cells mediate superior antitumor activity. JCI Insight 3, (2018).

